# Regulation of checkpoint kinase signalling and tumorigenesis by the NF-κB regulated gene, CLSPN

**DOI:** 10.1101/358291

**Authors:** Jill E. Hunter, Jacqueline A. Butterworth, Helene Sellier, Saimir Luli, Achilleas Floudas, Adam J. Moore, Huw D. Thomas, Kirsteen J. Campbell, Niall S. Kenneth, Robson T. Chiremba, Dina Tiniakos, Andrew M. Knight, Benjamin E. Gewurz, Fiona Oakley, Michelle D. Garrett, Ian Collins, Neil D. Perkins

## Abstract

Inhibition of the tumour promoting activities of NF-κB by cell signalling pathways has been proposed as a natural mechanism to limit the development of cancer. However, there has been a lack of evidence for these effects *in vivo*. Here we report that *RelA*^T505A^ mice, where a CHK1 targeted Thr505 phosphosite is mutated to alanine, display earlier onset of MYC driven lymphoma than wild type littermates. We describe a positive feedback loop in which the NF-κB subunits RelA and c-Rel, in a manner dependent upon RelA Thr505 phosphorylation, drive the expression of the ATR checkpoint kinase regulator Claspin in response to DNA replication stress in cancer cells. This in turn is required for maintenance of CHK1 activity. Loss of a single allele of the *Clspn* gene in mice is sufficient to drive earlier tumorigenesis and low levels of *CLSPN* mRNA expression are associated with worse survival in some forms of human cancer. We propose that loss of this pathway early in tumorigenesis promotes cancer development through increased genomic instability. However, in malignant cancer cells it can help promote their addiction to the checkpoint kinase signalling required for the maintenance of genomic integrity. Importantly, disruption of this pathway leads to resistance of cells to treatment with CHK1 inhibitors. Claspin expression could therefore act as a biomarker for responsiveness of patients to CHK1 inhibitors and provide a potential pathway for the development of tumour resistance.

## Introduction

The Nuclear Factor κB (NF-κB) family of transcription factors, comprising RelA/p65, RelB, c-Rel, p50/p105 (NF-κB1) and p52/p100 (NF-κB2), are important regulators of cancer cell biology (Perkins, 2012). Through their ability to regulate a wide variety of genes associated with inflammation, proliferation, apoptosis and metastasis, aberrant NF-κB subunit activity can promote the growth, survival and spread of tumour cells (Perkins, 2012). However, many *in vivo* studies have relied upon mutation or deletion of subunits of the IκB Kinase (IKK) complex, which drives activation of the NF-κB pathway but has also been reported to have many other functions (Hinz and Scheidereit, 2014). By contrast, the functions of specific NF-κB subunits have often not been explored. Indeed, while there is often an assumption that NF-κB is an obligate tumour promoter, tumour suppressor like characteristics have been identified *in vitro* that are rarely examined using *in vivo* models (Perkins, 2012). For example in cell lines, in response to inducers of DNA replication stress, NF-κB can have a pro-apoptotic function (Campbell et al., 2006; Msaki et al., 2011; Rocha et al., 2005; Wu and Miyamoto, 2008).

Given the many diverse roles of the NF-κB pathway, dissecting the complexity of its role in different contexts and diseases can be challenging. In particular, mouse gene knockouts of NF-κB will remove many functions simultaneously and are thus a crude tool to analyse the contribution of different, sometimes apparently opposing activities. By contrast, knockin mutations of NF-κB subunits in mice that inactivate, for example, specific sites of post-translational modifications, can potentially reveal aspects of their behaviour not seen with knockouts, inhibition through RNA interference, or expression of the inhibitor IκBα (Riedlinger et al., 2018). An example of this is the *RelA*^T505A^ mouse (Moles et al., 2016). Previously this laboratory has shown that in cell lines, CHK1 can phosphorylate the RelA C-terminal transactivation domain at Thr505 (T505), resulting in inhibition of tumour promoting activities of NF-κB, including resistance to apoptosis, autophagy and cell proliferation/migration (Campbell et al., 2006; Msaki et al., 2011; Rocha et al., 2003; Rocha et al., 2005). However, the *in vivo* significance of these effects was not known. Therefore to learn more about this we created a knock-in mouse model in which the RelA Thr 505 residue (504 in the mouse) is mutated to Ala (T505A). Importantly, and consistent with our model for the function of RelA T505 phosphorylation, we previously reported that this mutation led to earlier onset of hepatocellular carcinoma (HCC) *in vivo* using the N-nitrosodiethylamine (DEN) model (Moles et al., 2016). However, the mechanisms leading to this effect *in vivo* were not clear. Therefore, in this study, to better understand the basis of why inactivating RelA T505 phosphorylation leads to earlier onset of cancer, we decided to investigate this pathway in the context of oncogene driven DNA replication stress. We chose the Eµ-Myc mouse model of B-cell lymphoma to perform these experiments since it is a well-established and highly characterised system (Harris et al., 1988). Moreover, over-expression of the oncogene c-Myc is a feature of many types of cancer and results in DNA replication stress leading to genomic instability and tumorigenesis (Dominguez-Sola and Gautier, 2014; Maya-Mendoza et al., 2015; Rohban and Campaner, 2015). Since DNA replication stress leads to activation of the checkpoint kinases Ataxia Telangiectasia and Rad3 Related (ATR) and CHK1, given our previous observations, this had the potential to be an ideal model system to further explore the functional role of RelA T505 phosphorylation.

Here we report that mutation of RelA at T505 also leads to earlier onset of lymphoma in the Eµ-Myc mouse. Importantly, we identify the regulator of ATR/CHK1 signalling CLSPN (Claspin) gene as an important effector of both the RelA and c-Rel NF-κB subunits in response to DNA replication stress in cancer. In *RelA*^T505A^ and *c-Rel* null mice, we demonstrate reduced levels of CLSPN expression, leading to reduced levels of CHK1 activity. This in turn, leads to resistance of these cells to treatment with a highly selective and specific CHK1 inhibitor, a result with important implications for the use of such compounds in the clinic. Moreover, for the first time, we demonstrate that loss of a single allele of CLSPN is sufficient to drive early tumorigenesis in mice and that low CLSPN transcript levels are associated with significantly reduced survival of human patients with certain types of cancer.

## Results

### RelA T505 phosphorylation is required for pro-apoptotic effects of NF-κB following DNA replication stress

RelA Thr 505 (T505) phosphorylation has been shown to induce a pro-apoptotic form of NF-κB following stimulation with the chemotherapeutic drug and DNA cross-linker cisplatin (Campbell et al., 2006; Msaki et al., 2011). However, these experiments were based on exogenous expression of wild type (WT) and T505A RelA mutants in either human cell lines or *Rela*^-/-^ immortalised mouse embryonic fibroblasts (MEFs). Therefore, to confirm the role of endogenous RelA T505 phosphorylation as a regulator of the NF-κB response to DNA damage we generated immortalised and primary MEFs from *RelA*^T505A^ mice and matching WT littermates. Importantly, and in agreement with the previous data, mutation of endogenous RelA at T505 strongly protected cells from cisplatin-induced apoptosis (Fig. 1A & B, S1A). Induction of p53 protein in response to cisplatin treatment was not significantly affected in early passage *RelA*^T505A^ immortalised MEFs (Fig. 1A). Western blot analysis revealed no significant effects on the other NF-κB subunits in the *RelA*^T505A^ MEFs (Fig S1B).

**Figure 1.**
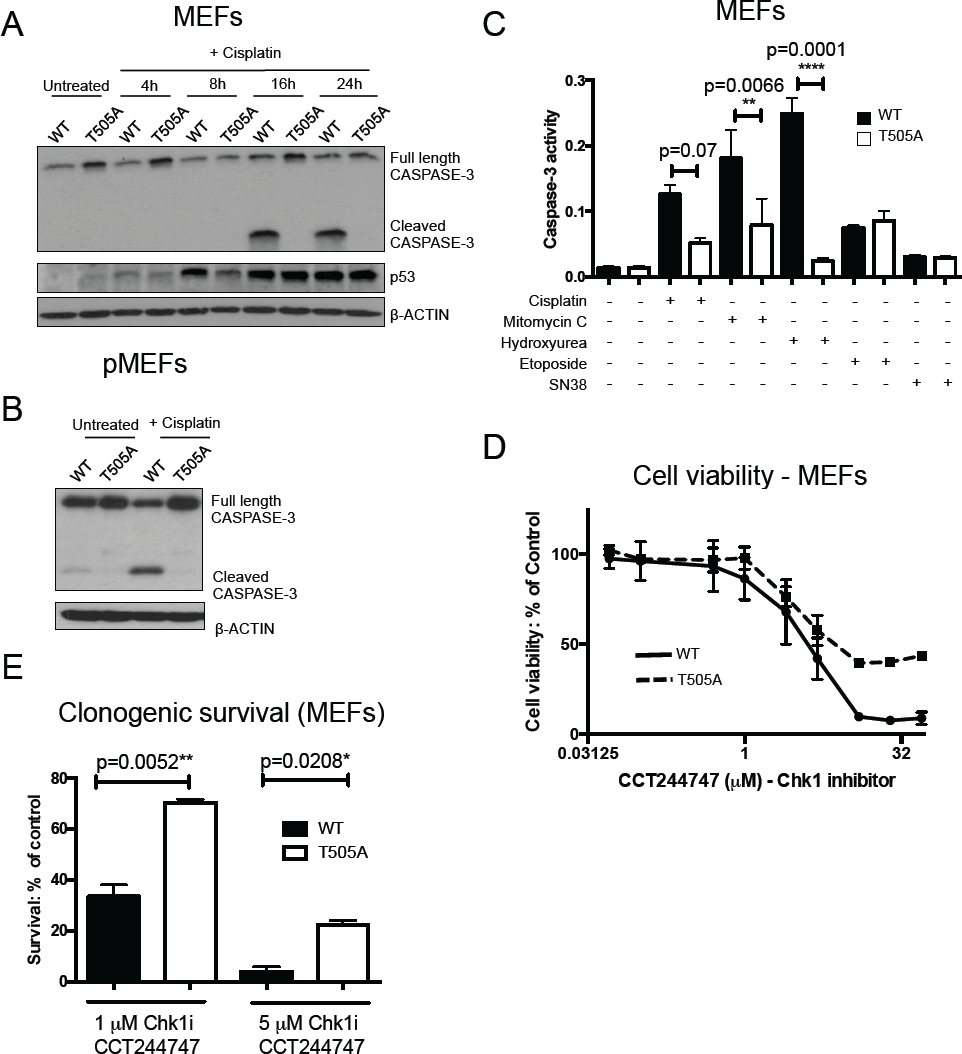
RelA T505 phosphorylation is required for pro-apoptotic effects of NF-κB following DNA replication stress. (A & B) *RelA*^T505A^ MEFs are resistant to Cisplatin induced apoptosis. Western blot analysis of full length and cleaved CASPASE 3 in (A) immortalised or (B) primary wild type (WT) and RelA T505A MEFs after treatment with the DNA damaging agent, Cisplatin (4 μg/ml). (C) *RelA*^T505A^ MEFs are resistant to apoptosis resulting from treatment with inducers of DNA replication stress. CASPASE 3 activity assay in immortalised WT and *RelA*^T505A^ MEFs after treatment with Cisplatin (4 μg/ml), Mitomycin C (1 μg/ml)., Hydroxyurea (0.5 mM), Etoposide (15 μM) and the active metabolite of Camptothecin, SN38 (5 μM). All drugs were added for 16 hours before analysis apart from SN38 (48 hours). Results shown are the mean + SEM from 3 separate repeat experiments. (D) *RelA*^T505A^ MEFs are resistant to CHK1 inhibitor treatment. Cell viability (Prestoblue assay) in WT and *RelA*^T505A^ MEFs following treatment with increasing concentrations of the CHK1 inhibitor, CCT244747 for 72 hours. (E) Increased clonogenic survival in *RelA*^T505A^ MEFs following CHK1 inhibitor treatment Clonogenic survival in WT and *RelA*^T505A^ MEFs following either treatment with either 1 μM (p=0.0052 ** Unpaired Student’s t-test) or 5 μM (p=0.0208 * Unpaired Student’s t-test) of the CHK1 inhibitor, CCT244747 for 24 hours.

We extended this analysis to other inducers of DNA damage and similarly the RelA T505A MEFs again displayed less apoptosis in response to the DNA crosslinker mitomycin C and hydroxyurea, an inhibitor of ribonucleotide reductase, both of which are inducers of DNA replication stress (Fig. 1C). By contrast, there was no significant difference in apoptosis between WT and *RelA*^T505A^ MEFs when treated with etoposide, a Topoisomerase II inhibitor, or with SN38, a Topoisomerase I inhibitor and the active metabolite of camptothecin (Fig. 1C).

Since the RelA T505 residue can be phosphorylated by the checkpoint kinase CHK1 following replication stress (Rocha et al., 2005), we were interested in whether the RelA T505A mutation would also affect cell survival in response to treatment with CHK1 inhibitors, an important class of novel cancer therapeutics currently in clinical trials (Carrassa and Damia, 2017). Interestingly, in both cell viability and clonogenic survival assays the *RelA*^T505A^ MEFs were significantly more resistant to the CHK1 inhibitor CCT244747 (Fig. 1D & E)(Walton et al., 2012), and the structurally unrelated CHK1 inhibitor MK8776 (Fig. S1C) (Daud et al., 2015; Guzi et al., 2011) compared with matched WTs.

Taken together these results confirm and extend the original observation that phosphorylation at the RelA T505 residue is an important regulator of the cellular response to inducers of DNA replication stress. Furthermore, they demonstrate that the *RelA*^T505A^ mouse is a suitable model to interrogate the function of NF-κB in this context *in vivo* in a manner that cannot be performed with *RelA* knockout mice or other NF-κB pathway mutants.

### Mutation of RelA T505 results in earlier onset of Eµ-Myc driven lymphoma

Given our data supporting a critical role for RelA T505 phosphorylation as a regulator of how cells respond to DNA replication stress we decided to investigate its function in the context of c-Myc driven cancer. Over-expression of c-Myc is a feature of many types of cancer and results in DNA replication stress leading to genomic instability and tumorigenesis (Dominguez-Sola and Gautier, 2014). We chose the Eµ-Myc mouse model of B-cell lymphoma to perform these experiments since it is a well-established and highly characterised system (Harris et al., 1988).

Based on our data in Fig 1, our prediction was that the RelA T505A mutation should lead to earlier onset of lymphomagenesis and indeed homozygous Eµ-Myc*/Rela^T505A^* mice had a significantly shorter overall survival (median survival 83.5 days) when compared with the Eµ-Myc mice (median survival 122 days) (Fig. 2A). Moreover, consistent with RelA T505 phosphorylation being a regulator of the response to DNA replication stress, we observed significantly higher levels of γH2AX, a marker of DNA damage and genomic instability, in Eµ-Myc*/Rela^T505A^* lymphomas (Figs. 2B-D).

**Figure 2.**
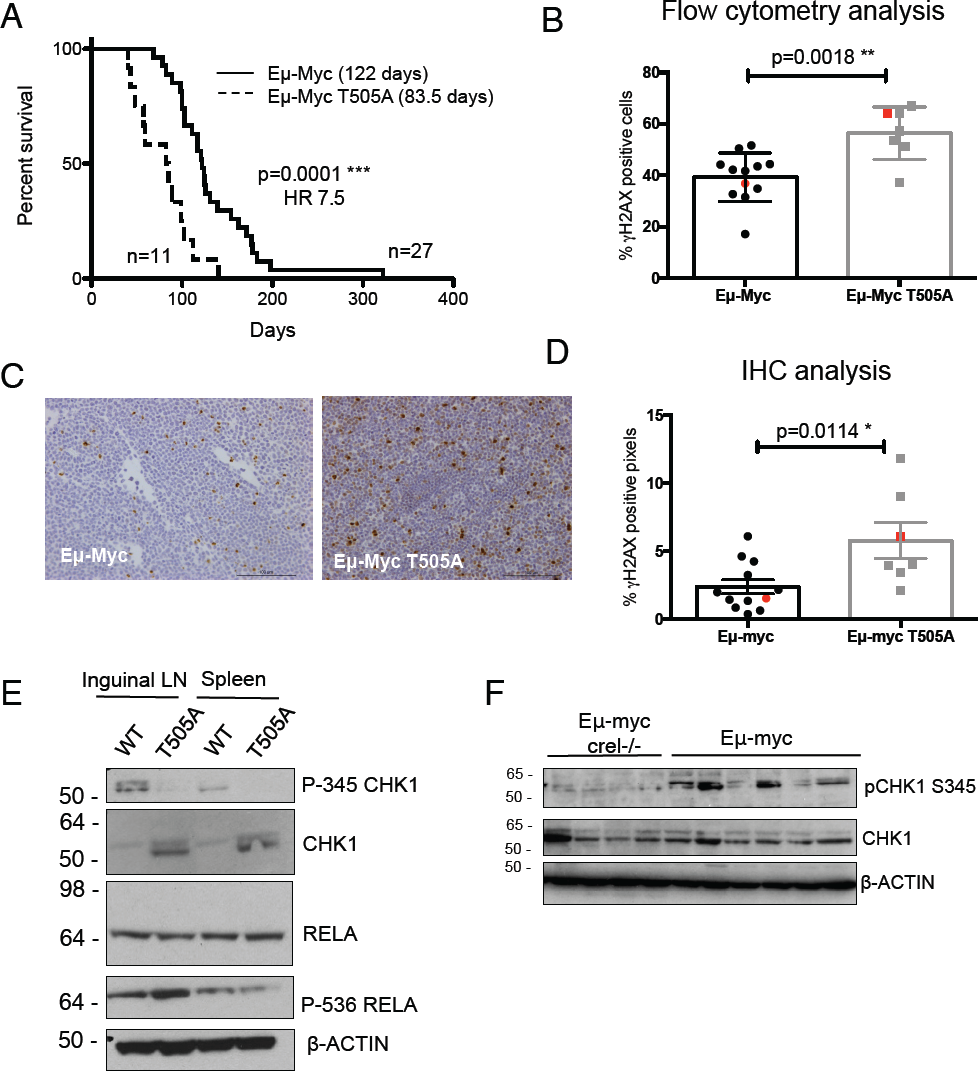
The RelA T505A mutation leads to earlier onset of lymphoma, genomic instability and a defect in CHK1 activation. (A) Earlier inset of lymphoma in Eµ-Myc*/Rela^T505A^* mice. Kaplan Meier survival curves for Eµ-Myc (n=27) and homozygous Eµ-Myc*/Rela^T505A^* male mice (n=11). Survival is significantly shorter in Eµ-Myc/*RelA*^T505A^ mice (p=0.0001 *** Mantel-Cox test) and hazard ratio (HR) analysis indicates that these mice are at 7.5 times greater risk of dying earlier due to lymphoma compared with Eµ-Myc mice. The median survival for each genotype is indicated. (B-D) Increased genomic instability in lymphomas from Eµ-Myc*/Rela^T505A^* mice. Flow cytometric analysis (B) of B cells prepared from Eµ-Myc and Eµ-Myc*/Rela^T505A^* spleens and stained for γH2AX, a marker of DNA damage (p=0.0018 ** Unpaired Student’s t-test). (C) Representative IHC images of Eµ-Myc and Eµ-Myc/*Rela^T505A^* lymph nodes stained with an antibody against γH2AX and counterstained with heamatoxylin. Brown staining indicates cells positive for γH2AX. Quantification of γH2AX positive pixels by IHC analysis (D) in Eµ-Myc and Eµ-Myc/*Rela^T505A^* lymph nodes. Each dot represents one mouse and at least blinded 5 fields of view were analysed per mouse (p=0.0114 * Unpaired Student’s t-test). The red dot in each case illustrates the mouse shown in (C). (E & F) *Eµ-Myc/Rela^T505A^* lymphomas have a defect in CHK1 activation. Western blot analysis of phospho-S345 CHK1, CHK1, RELA and phospho-S536 RELA in snap frozen tumour extracts prepared from Eµ-Myc and *Eµ-Myc/Rela^T505A^* mouse inguinal lymph nodes and spleens. Red dots in scatter plots in (B) and (D) indicate the mice used for protein analysis. (F) Eµ-Myc*/c-Rel^-/-^* lymphomas also have a defect in CHK1 activation. Western blot analysis of phospho-S345 CHK1 and CHK1using extracts prepared from Eµ-Myc and Eµ-Myc*/c-Rel^-/-^* tumorigenic spleens.

Further analysis of the lymphoma cells from Eµ-Myc*/Rela^T505A^* mice did not reveal any significant differences in the expression of NF-κB anti-apoptotic target gene *Bcl-xL* (*Bcl2l1*) (Fig S2A). Nor did we observe differences in expression of the pro-apoptotic gene *Bax* between WT Eµ-Myc and Eµ-Myc*/Rela^T505A^* lymphomas (Fig. S2B). Furthermore, flow cytometric analysis revealed that the percentages of B and T lymphocyte sub-populations, along with macrophage and dendritic cell populations, were similar in WT and *RelA*^T505A^ mice, indicating that a fundamental difference in the immune system of these mice was unlikely to be the cause of the earlier onset of lymphoma observed in Eµ-Myc*/Rela^T505A^* mice (Fig. S2C and Suppl Table 1). Western blot analysis showed no significant effects on RelA and c-Rel levels in Eµ-Myc*/Rela^T505A^* cells, although apparently raised levels of RELB and reduced processing of p105 to p50 were observed (Fig. S2D).

### Eµ-Myc*/Rela^T505A^* and Eµ-Myc*/c-Rel^-/-^* lymphomas exhibit a defect in CHK1 signalling

Development of lymphoma in the Eµ-Myc model requires DNA mutations leading to, for example, loss of function of the ARF/p53 tumour suppressor pathway (Eischen et al., 1999). This suggested that the increased genomic instability seen in the Eµ-Myc*/Rela^T505A^* lymphomas could provide a potential explanation for the earlier lymphoma onset observed. We therefore decided to examine the ATR/CHK1 DNA replication stress response pathway in more detail. Significantly, levels of CHK1 phosphorylated at Ser345, a marker for its activation by ATR, were reduced in Eµ-Myc*/Rela^T505A^* cells (Fig. 2E and see also 4C). These effects did not result from problems with extract preparation since analysis of the same extracts revealed unchanged or increased levels of phospho-ERK, JNK and p38 MAP kinase together with RELA phosphorylated at Ser536 (Fig. 2E and S2E). Unfortunately, our phospho-RELA T505 antibody does not work well with extracts from mouse tissues, meaning we could not confirm phosphorylation at this site.

In a parallel series of experiments we have also found that knockout of the c-Rel NF-κB subunit results in earlier lymphoma onset in the Eµ-Myc model (Hunter et al., 2016). We were therefore interested in whether Eµ-Myc/*c-Rel*^-/-^ lymphoma cells would also display loss of CHK1 activity. Interestingly, western blot analysis also demonstrated a lower level of Ser345 phosphorylated CHK1 in Eµ-Myc*/c-Rel^-/-^* cells (Fig. 2F). These results indicated that, surprisingly, Eµ-Myc*/Rela^T505A^* and Eµ-Myc*/c-Rel^-/-^* lymphomas share a common and intrinsic defect in CHK1 kinase signalling.

### Eµ-Myc*/c-Rel^-/-^* and Eµ-Myc/*Rela*^T505A^ lymphoma cells are resistant to CHK1 inhibition

Disruption of the DNA damage response, frequently seen as a consequence of genetic mutation, can lead to more rapid onset of tumorigenesis (Campaner and Amati, 2012). However, inhibiting those pathways that are still intact provides a potential therapeutic strategy for targeting tumours that have become dependent on their activity (Garrett and Collins, 2011). For this reason, inhibitors of ATR and CHK1 represent a potential new class of anti-cancer therapies, and are currently in clinical trials (Carrassa and Damia, 2017).

Given the apparent defect in CHK1 kinase signalling in Eµ-Myc*/Rela^T505A^* and Eµ-Myc*/c-Rel^-/-^* lymphomas, we hypothesised that a consequence would be altered sensitivity to CHK1 inhibition *in vivo*, similar to the increased resistance we had seen in *RelA*^T505A^ MEFs (Figs 1D & E, S2C). The CHK1 inhibitor CCT244747 has shown efficacy in a transgenic model of MYCN-driven neuroblastoma (Walton et al., 2012). Moreover, we have previously shown that the related CHK1 inhibitor CCT245737 inhibits the growth of re-implanted WT Eµ-Myc cells (Walton et al., 2015).

We first examined how WT Eµ-Myc, Eµ-Myc/*c-Rel^-/-^* or Eµ-Myc/*Rela*^T505A^ lymphoma cells responded to treatment with CCT244747 for 96 hours *ex vivo.* Consistent with this hypothesis, we observed small but significant differences, with WT cells having reduced viability relative to Eµ-Myc*/c-Rel^-/-^* or Eµ-Myc/*Rela*^T505A^ tumour cells (Fig. 3A). We next evaluated the effectiveness of CCT244747 *in vivo* by analysing its effect on the growth of five transplanted WT Eµ-Myc, Eµ-Myc/*c-Rel^-/-^* and Eµ-Myc/*Rela*^T505A^ tumours. Each tumour was implanted into six syngeneic C57Bl/6 recipient mice and three were treated orally with CCT244747 once a day for nine days, while three received a vehicle control (implantation and dosing schematic shown in Fig. 3B). After the 9-day course of treatment, we observed a striking reduction in lymphoid tumour burden in all mice re-implanted with WT Eµ-Myc tumours (Figs. 3C & D, S3A-F). Some splenomegaly was observed in mice treated with CCT244747 (Fig. 3D) but other, non-lymphoid organs were unaffected by the drug (not shown). By contrast, both Eµ-Myc*/c-Rel^-/-^* and Eµ-Myc/*Rela*^T505A^ transplanted tumours were largely resistant to CCT244747 treatment (Figs. 3C & D, S3A-F), consistent with the *ex vivo* results (Fig. 3A). All of the Eµ-Myc/*Rela*^T505A^ tumours and four of the five Eµ-Myc*/c-Rel^-/-^* tumours showed no significant reduction in lymphoid tumour burden after CCT244747 treatment, whereas one tumour exhibited a partial response with a reduction in tumours of the thymus and cervical lymph nodes. These data confirmed that the NF-κB mutant Eµ-Myc tumours have disrupted checkpoint kinase signalling pathway, bypassing a requirement for CHK1 activation, thus rendering them insensitive to the CHK1 inhibitor.

**Figure 3.**
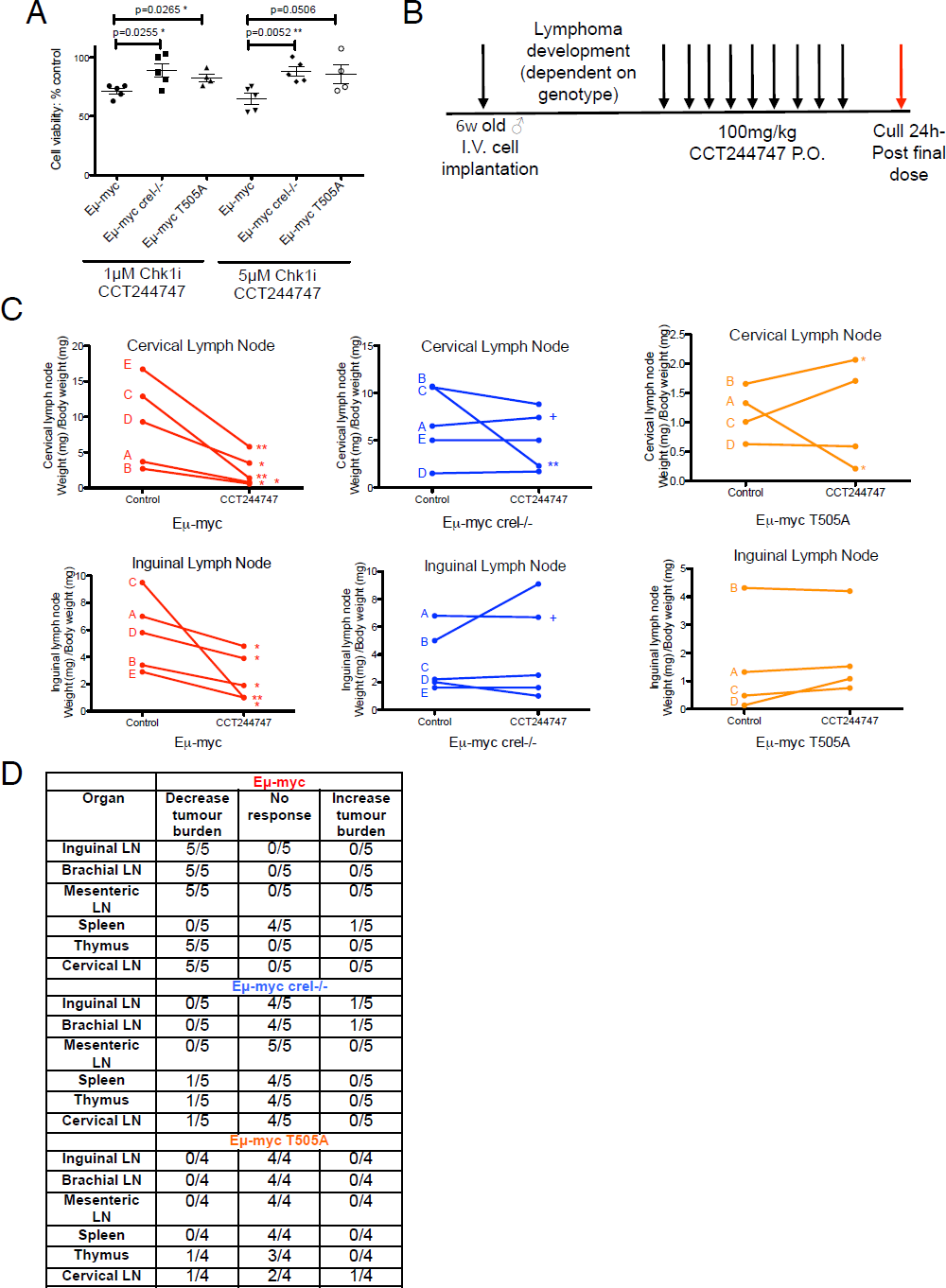
Eµ-Myc/*Rela^T505A^* and Eµ-Myc*/c-Rel^-/-^* lymphomas are resistant to CHK1 inhibition (A) Eµ-Myc/*c-Rel^-/-^ and* Eµ-Myc/*Rela^T505A^* lymphoma cells are resistant to CHK1 inhibition *ex vivo*. Eµ-Myc, Eµ-Myc/*c-Rel^-/-^ and* Eµ-Myc/*Rela^T505A^* lymphoma cells were treated with 0.5µM or 1µM CCT244747 (or vehicle control) for 96 hours *ex vivo*. WT Eµ-Myc tumour cells show a reduced cell viability compared with the Eµ-Myc/*c-Rel^-/-^* or Eµ-Myc/ *RelA*^T505A^ tumour cells at all doses of CCT244747 tested. (B) Schematic of the *in vivo* study using the CHK1 inhibitor, CCT244747. 5 lymphomas of each genotype were reimplanted separately into 6 syngeneic recipient C57Bl/6 mice. Lymphomas were allowed to develop and when palpable 3 were treated with CCT244747 (100 mg/kg p.o), and 3 with vehicle control, for 9 days. (C) Reimplanted Eµ-Myc/*c-Rel^-/-^ and* Eµ-Myc/*Rela^T505A^* lymphoma cells are resistant to CHK1 inhibition *in vivo.* Line graphs showing the mean response of the five reimplanted Eµ-Myc (red) and Eµ-Myc/*c-Rel-/-* (blue) and Eµ-Myc/*Rela^T505A^* (orange) tumours and their response to CCT244747. A response was defined as a significant reduction (or increase) in tumour burden (p<0.05) using unpaired Student’s t-tests. The complete data set is summarised in (D). ‘+’ indicates one experiment where treatment was stopped after 7 days and the mice were humanely killed at this timepoint due to the disease burden in these mice. (D) Table showing the response of 5 re-implanted Eµ-Myc, Eµ-Myc/*c-Rel^-/-^* and Eµ-Myc/*Rela^T505A^* tumours to CCT244747, in all sites where lymphoid tumour burden is anticipated in this model.

### Loss of Claspin expression in Eµ-Myc*/Rela^T505A^* and Eµ-Myc*/c-Rel^-/-^* lymphomas

We next investigated the mechanistic basis of the loss of CHK1 activity in the NF-κB mutated cells. A number of proteins are known to regulate the DNA replication stress checkpoint response to oncogene activation, which includes the coregulators ATRIP, Claspin (encoded by the *Clspn* gene in mice), TopBP1 and Rad17 that are all required for ATR mediated activation of CHK1 (Fig 4A) (Errico and Costanzo, 2012). qRT-PCR analysis of RNA extracted from Eµ-Myc lymphoma cells revealed that of these genes, only *Clspn* expression was affected by loss of c-Rel or the RelA T505A mutation (Fig. 4B and S4A-E). By contrast, *Clspn* levels were similar in normal B-cells from WT, *RelA*^T505A^ and *c-Rel^-/-^* mice (Fig. S5A & B) suggesting that it is a NF-κB target gene only under certain conditions, such as oncogene (e.g. MYC) induced DNA replication stress. Claspin is an adaptor protein associated with DNA replication forks that is required for ATR dependent phosphorylation of CHK1 following DNA replication stress and other forms of DNA damage. Claspin can also function as a sensor to monitor the integrity of DNA replication forks through binding to branched DNA structures (Azenha et al., 2017; Chini and Chen, 2004; Errico and Costanzo, 2012; Kumagai and Dunphy, 2003). Claspin inactivation contributes to malignant transformation of cell lines by oncogenic viruses (Hein et al., 2009; Koganti et al., 2014; Spardy et al., 2009) such as human papillomavirus. HPV-E7 protein can induce Claspin degradation and thus bypass cellular checkpoints allowing the virus to replicate in the presence of DNA damage, (Spardy et al., 2009). Proteolytic degradation of Claspin has also been shown to disrupt ATR/CHK1 signalling and promote EBV transformation of primary B-cells (Koganti et al., 2014). Consequently, it has been suggested that Claspin inactivation could be an essential event during carcinogenesis (Azenha et al., 2017), although evidence for this at the genetic level has not been established. Moreover, Rad17, a vital component of the replication stress response has previously been reported as a putative tumour suppressor in murine lymphoma (Bric et al., 2009), indicating that this pathway is essential for the maintenance of genome integrity.

**Figure 4.**
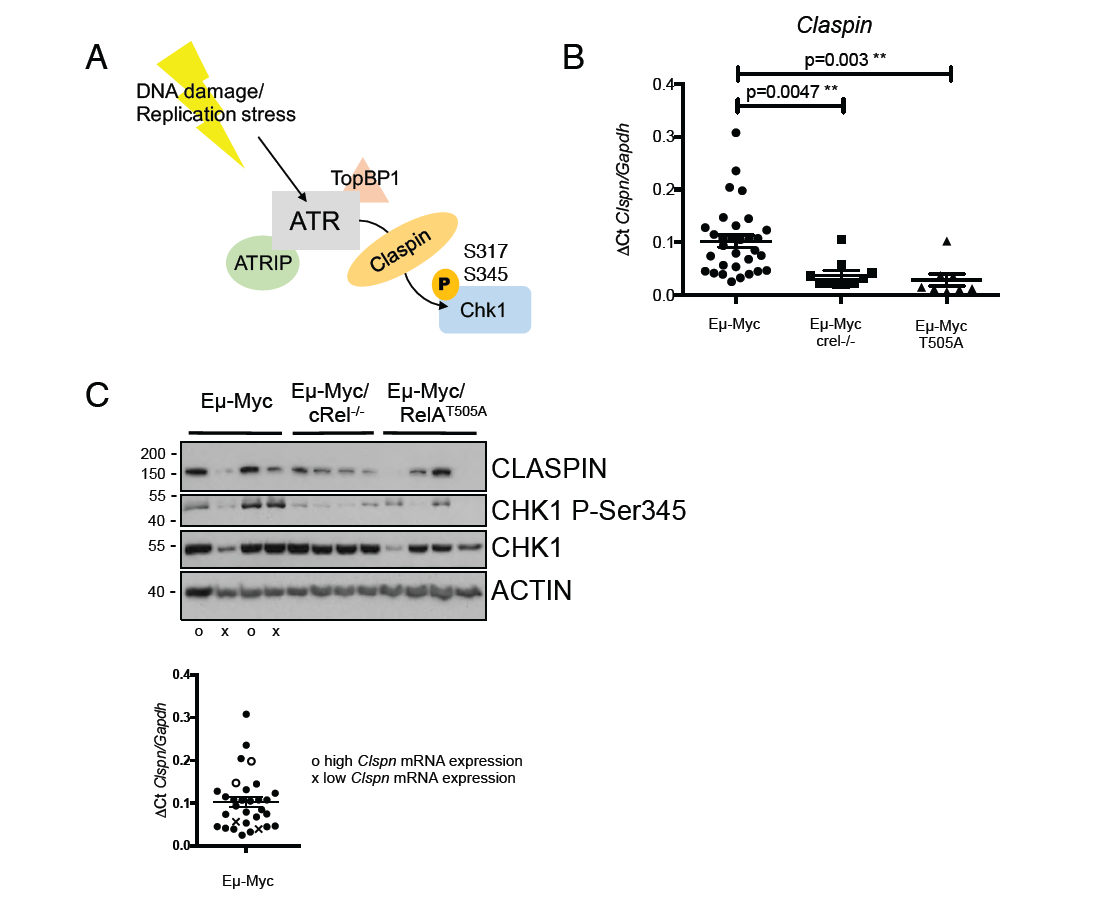
Eµ-Myc*/c-Rel^-/-^* and Eµ-Myc/*RelA*^T505A^ lymphomas fail to induce Claspin expression. (A) Schematic diagram illustrating the activation of CHK1 by ATR following DNA damage or replication stress, and the role of Claspin as an essential adaptor protein. (B) Reduced Clspn mRNA expression in Eµ-Myc*/c-Rel^-/-^* and Eµ-Myc/*Rela^T505A^* lymphomas. qRT-PCR data showing relative *Clspn* mRNA expression in tumorigenic spleens from Eµ-Myc (n=30), Eµ-Myc/*c-Rel^-/-^* (n=11) and Eµ-Myc/*Rela^T505A^* (n=8) mice. Data represents mean ± SEM. p** <0.01 (Unpaired Student’s t-test), each point is an individual mouse. (C) Claspin protein expression is also reduced in Eµ-Myc*/c-Rel^-/-^* and Eµ-Myc/*Rela^T505A^* lymphomas. Western blot analysis of Claspin using extracts prepared from Eµ-Myc, Eµ-Myc*/c-Rel^-/-^* or Eµ-Myc/*Rela^T505A^* tumorigenic spleens. Accompanying scatter plot illustrates which wild type mice with either high (o) or low (x) *Clspn* mRNA expression were used for protein analysis.

In human cell lines derived from solid tumours, the *CLSPN* gene has previously been shown to be a direct c-Rel target gene (Kenneth et al., 2010). To extend this analysis we extracted data from a published ChIP-seq analysis of the EBV-transformed human lymphoblastoid B-cell line (LCL) GM12878 (Zhao et al., 2014). This demonstrated that the *CLSPN* promoter region can potentially be a target of all NF-κB subunits in B-cells, including RelA and c-Rel (Fig. S5C).

Western blot analysis confirmed that there was a correlation between low *Clspn* mRNA levels and low Claspin protein levels in WT Eµ-Myc cells (Fig. 4C). Moreover Claspin protein levels were reduced in both Eµ-Myc*/Rela^T505A^* and Eµ-Myc*/crel^-/-^* lymphoma cells, although one Eµ-Myc*/Rela^T505A^* sample displayed aberrantly high levels of protein despite low mRNA levels (Fig. 4C). Claspin protein levels are also regulated by proteolysis and this result suggests that some tumour cells overcome the reduction of mRNA expression through increased protein stability (Koganti et al., 2014; Yuan et al., 2014; Zhu et al., 2014). NF-κB regulation of *Clspn* gene expression therefore provides one potential explanation for the defects in CHK1 activity we observed in Eµ-Myc*/Rela^T505A^* and Eµ-Myc*/c-Rel^-/-^* lymphomas.

### Claspin levels predict outcome in Eµ-Myc lymphoma

In our analysis of CHK1 phosphorylation we noticed that this was highly variable in the WTEµ-Myc tumour cells (Fig. 2F & 4C). Moreover, *Clspn* mRNA expression also varied in WT Eµ-Myc cells, with a population showing very low levels of expression similar to that seen in Eµ-Myc*/Rela^T505A^* and Eµ-Myc*/c-Rel^-/-^* cells (Fig. 4B). We concluded that if low levels of Claspin and CHK1 activity in the *c-Rel*^-/-^ and *Rela*^T505A^ cells were leading to earlier onset of disease then this would also be seen in this population of WT Eµ-Myc cells with low *Clspn* mRNA levels. To test this hypothesis, we analysed the survival of the two populations of Eµ-Myc mice identified with *Clspn* mRNA expression either below or above the median level (Fig. S5D). Importantly, this revealed that low levels of this transcript were associated with significantly earlier onset of lymphoma (median onset 82 versus 138 days) (Fig. 5A). This was almost identical to that found in Eµ-Myc mice lacking c-Rel (median 79 days) or to that seen in *Rela*^T505A^ mutant mice (median 83.5 days). By contrast, analysis of below and above the median mRNA levels of *Atr*, *Atrip*, *Topbp1*, *Rad17* and *Chek1* showed no correlation with the survival times of Eµ-Myc mice (Fig. S4A-E).

**Figure 5.**
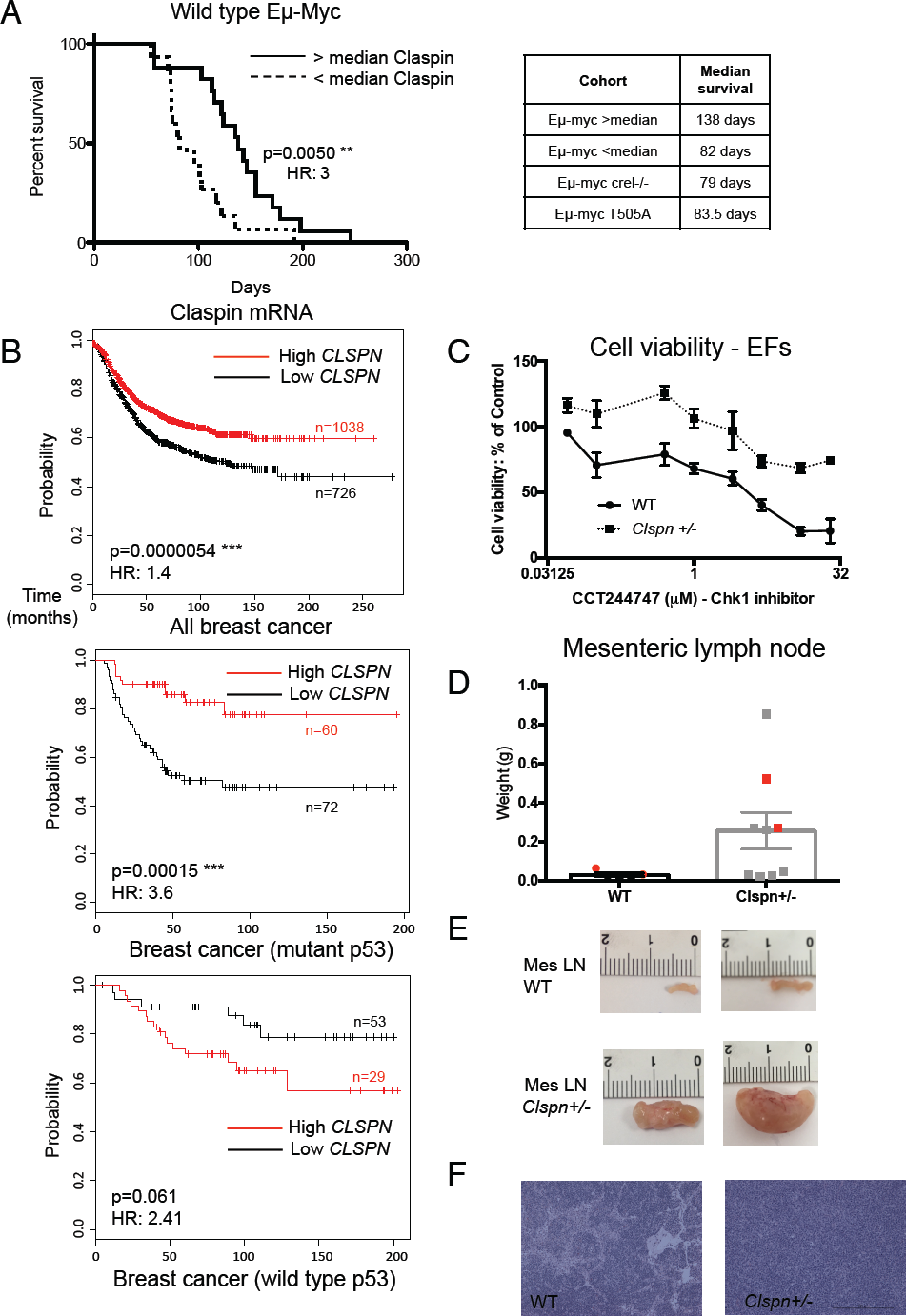
Low Claspin levels are associated with poor outcome in Eµ-Myc lymphomas, human breast cancers and lead to spontaneous B-cell lymphoma in aged *Clspn*^+/-^ mice. (A) WT Eµ-Myc mice with lower Claspin levels develop lymphoma earlier. Kaplan-Meier survival analysis of WT Eµ-Myc mice comparing above and below median level expression of *Clspn* mRNA. Table detailing the survival of Eµ-Myc and Eµ-Myc/*c-Rel^-/-^* mice as described previously in (Hunter et al., 2016). (B) Low *CLSPN* mRNA levels are associated with poor survival in breast cancer patients with mutant p53. Kaplan-Meier survival analysis of breast cancer datasets from KM Plotter with differing levels of *CLSPN* mRNA (auto select best cutoff). Patients with low *CLSPN* mRNA levels have significantly shorter overall survival (p=0.0000054 Mantel-Cox test, HR 1.4) and this is consistent with the results in the Eµ-Myc mouse model. In breast cancer patients with mutant p53 mutant breast cancer low *CLSPN* levels also have significantly shorter overall survival (p=0.00015 Mantel-Cox test, HR 3.6), while the situation is reversed in breast cancer patients with WT p53 (p=0.0033 Mantel-Cox test, HR 2.6). (C) Primary fibroblasts from *Clspn*^+/-^ mice are resistant to CHK1 inhibition. Cell viability (Prestoblue assay) in WT and *Clspn*^+/-^ primary ear fibroblasts (EFs) following treatment with increasing concentrations of the CHK1 inhibitor, CCT244747. (D – F) 18 month old *Clspn*^+/-^ spontaneously develop B-cell lymphoma. (D) Weight of the mesenteric lymph nodes from 18 month old WT and CLSPN^+/-^ mice. (E) Representative images of the mesenteric lymph node from two 18 month old WT and two 18 month old WT *Clspn*^+/-^ mice. (F) Representative H & E stained sections of the mesenteric lymph node from 18 month old WT and *Clspn*^+/-^ mice. Red dots in scatter plot in (D) indicate samples used in (E) and (F).

### Claspin is a novel prognostic indicator in breast and gastric cancer

This data suggested that a failure to induce Claspin expression in tumour cells leads to a defective oncogene-induced DNA damage response and increased genomic instability, which will contribute to the earlier onset of disease. We therefore next investigated whether *CLSPN* mRNA levels might also predict progression of human cancer. Analysis of some previous gene expression studies of human B-cell lymphomas (e.g. (Hummel et al., 2006)) is complicated by an older Affymetrix microarray probe (219621_at) being designed to an incorrect Genbank entry for the sequence in the 3’ untranslated region of the human *CLSPN* mRNA. However, using more recent publicly available datasets on KM Plotter (Lanczky et al., 2016) and Human Protein Atlas (proteinatlas.org), we were able to assess the prognostic implications of Claspin expression in many different types of human cancer (summarised in Table S2). Strikingly, analysis of breast, stomach, head & neck, rectal and gastric cancer patients revealed that low levels of *CLSPN* transcripts were significantly associated with a poor prognosis in the disease (Fig. 5B, Table S2), just as we had observed in our mouse model (Fig. 5A). However, this was not the case in all tumour types and in some cases, such as lung, liver and pancreatic cancers, the opposite effect was observed (Table S2). An indication of why this might be the case came from further analysis of the breast cancer data. Here, we found that low *CLSPN mRNA* expression was also associated with poor survival in patients with mutant p53 but that in patients with WT p53 the opposite effect was seen (Fig. 5B). Similarly, with HER2 positive patients, low *CLSPN mRNA* expression again correlated with poor survival and this effect was lost in HER2 negative patients (Fig. S5E). This indicates a more complex situation where the consequences of high or low Claspin expression will depend on other mutation events in the tumour. Nonetheless, these data suggest that Claspin levels may be prognostic for some classes of cancer and importantly could influence the response to therapy, particularly of checkpoint kinase inhibitors, in human patients.

### Primary *Clspn*^+/-^ cells have a defect in the replication stress signalling pathway

Our data suggested that reduced Claspin levels could explain the results seen in Eµ-Myc*/Rela^T505A^* and Eµ-Myc*/c-Rel^-/-^* lymphoma cells. However, this data was largely correlative and we wished to directly test whether reduced Claspin levels alone can be sufficient to affect the response to CHK1 inhibition and enhance tumorigenesis. We therefore decided to examine these effects in *Clspn* knockout mice (purchased from the UC Davis KOMP Repository). In these studies heterozygous knockout *Clspn* mice were used since, similar to other mice with knockouts of genes associated with replication stress, such as *Atr^-^* (O’Driscoll, 2009) and *Chek1^-^* (Liu et al., 2000), homozygous loss of the *Clspn* gene is embryonic lethal at E10.5 (our observation and http://www.mousephenotype.org/data/genes/MGI:2445153). We generated independent isolates of *Clspn*^+/-^ and WT primary ear fibroblasts (EFs) and showed that Claspin protein levels were reduced in the *Clspn*^+/-^ cells, but that the protein levels of other pathway components, ATR and CHK1 were unaffected (Fig. S6A). Importantly, and consistent with our earlier data, *Clspn*^+/-^ EFs were more resistant to CHK1 inhibition, compared to their WT counterparts (Fig 5C and S6B).

### Aged *Clspn*^+/-^ mice develop spontaneous lymphoid tumours

To investigate whether reduced levels of *Claspin* would lead to lymphomagenesis, we aged a colony of *Clspn*^+/-^ mice. Strikingly, we observed tumours of the mesenteric lymph node in five of the nine *Clspn*^+/-^ mice we aged to 18 months of age (Fig. 5D – F), and in two of these mice there was further lymphoid disease burden as well as tumours of the small intestine/colon (Fig S6C). There were no overt signs of cancer, at either the macro- or microscopic level in any of the age-matched WT littermates (Fig 5D-F, S6C). Interestingly, the mesenteric lymph node appeared larger at all time points at which animals were harvested (Fig S6D & E) suggesting that the malignant cells home to this location and proliferate *in situ* thus resulting in the tumour burden seen in Fig 5D-F. No overall differences were seen with other lymph nodes (Figs S6F-H).

### *RelA*^T505A^ and *Clspn*^+/-^ mice are more susceptible to DEN induced acute liver injury

The data from aged *Clspn*^+/-^ mice suggested that reduced Claspin levels could lead to earlier onset of tumorigenesis in other contexts. However due to a number of ethical and technical problems associated with the difficulty of generating enough Eµ-Myc/*Clspn*^+/-^ mice to perform an adequately powered study (see Eµ-Myc mouse section of Methods), we decided investigate this further using the N-nitrosodiethylamine (DEN) induced model of hepatocellular carcinoma (Heindryckx et al., 2009), where we had already established an effect of the RelA T505A mutation (Moles et al., 2016). We first challenged both the *RelA*^T505A^ and the *Clspn*^+/-^ mice with an acute dose of DEN (Fig. 6A). DEN is a DNA alkylating agent and following acute administration, will promote hepatocyte death followed by compensatory proliferation. This compensatory proliferation following DEN-induced injury is critical for tumour formation in this model. We had previously observed that livers from *RelA*^T505A^ mice exhibited increased proliferation following acute DEN treatment (Moles et al.,2016). Given our data from the Eµ-Myc model, which indicated that mice with reduced Claspin levels would be more susceptible to genomic instability, we hypothesised that similar to *RelA*^T505A^ mice, *Clspn*^+/-^ mice would be more susceptible to acute liver injury induced by DEN.

**Figure 6:**
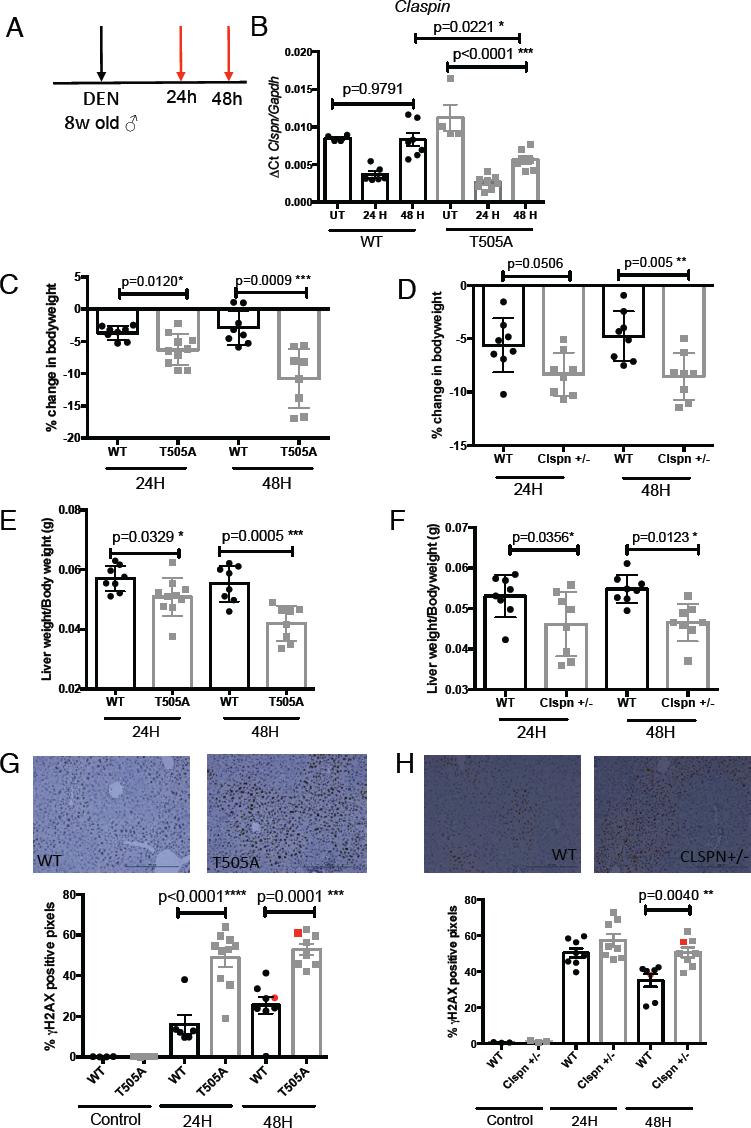
*RelA^T505A^* and *Clspn*^+/-^ mice are more susceptible to DEN induced acute liver injury. (A) Schematic diagram illustrating the acute DEN *in vivo* study in WT, *RelA*^T505A^ or *Clspn*^+/-^ mice. 80 mg/kg DEN in 0.9% saline was administered IP to 8 week old male mice. 24h or 48h post-DEN mice were humanely killed and tissues collected. (B) *Clspn* mRNA levels recover more slowly in *RelA*^T505A^ mice. qRT-PCR data showing relative *Clspn* mRNA expression in livers from WT and *RelA*^T505A^ mice 24h or 48h post DEN injury. Data represents mean ± SEM. p* <0.05 (Unpaired Student’s t-test). (C & D) *RelA*^T505A^ and *Clspn*^+/-^ mice show similar increased body weight loss after acute DEN treatment. Percentage change in bodyweight in WT, *RelA*^T505A^ (C) and *Clspn*^+/-^ (D) mice 24 h or 48 h post DEN injury. Data represents mean ± SEM. p* <0.05, p<0.01**, p<0.001*** (Unpaired Student’s t-test). (E & F) *RelA*^T505A^ and *Clspn*^+/-^ mice show similar increased liver damage after acute DEN treatment. Liver:body weight ratio at 24 h and 48 h post DEN injury in male WT, *RelA*^T505A^ (E) and *Clspn*^+/-^ (F). (G & H) *RelA*^T505A^ and *Clspn*^+/-^ mice both show increased levels of DNA damage after acute DEN treatment. Quantification of γH2AX positive pixels by IHC analysis in WT, *RelA*^T505A^ (G) and *Clspn*^+/-^ (H) livers either 24 h or 48 h post DEN injury. Each dot represents one mouse and at least blinded 5 fields of view were analysed per mouse. Data represents mean ± SEM. p<0.001*** p>0.0001**** (Unpaired Student’s t-test) Representative images are shown, with red dots in scatter plots indicating representative images chosen.

Following acute DEN administration, we observed a decrease in *Clspn* transcript levels after 24 hours, which in wild type mice returned to untreated levels after 48 hours. However, in the *RelA*^T505A^ mice, this recovery was significantly reduced (Fig. 6B).

Consistent with our hypothesis that *Clspn* transcript levels are regulated by RelA in a T505 dependent manner following DNA damage, acute DEN administration resulted in significant decreases in body weight of both the *RelA*^T505A^ (Fig 6C) and *Clspn*^+/-^ (Fig 6D) mice, compared with their respective WT counterparts, suggestive of greater damage and parenchyma loss. Consistent with this, the liver to bodyweight ratio in both *RelA*^T505A^ (Fig 6E) and *Clspn*^+/-^ (Fig 6F) mice at both time points post DEN administration was also significantly reduced. This decrease in the liver to bodyweight ratio is suggestive of increased hepatocyte damage, which was confirmed by the increased presence of γH2AX staining, a marker of DNA damage in *RelA*^T505A^ (Fig 6G) and *Clspn*^+/-^ (Fig 6H) compared to WT control mice following DEN. These results confirm that the reduction in Claspin expression in both the *RelA*^T505A^ and *Clspn*^+/-^ mice can have important consequences for how cells cope with DNA damage.

### Earlier onset of liver cancer in *Clspn*^+/-^ mice

We next investigated whether reduced Claspin levels would affect tumorigenesis in the chronic model of DEN-induced HCC. DEN was administered to 15-day-old WT and *Clspn*^+/-^ mice and tumour growth was evaluated after 30 weeks. At this point tumours start to become visible on the liver surface in WT mice and therefore, similar to our study using the *RelA*^T505A^ mice (Moles et al., 2016) this is the optimal time to investigate if disruption of the *Clspn* gene resulted in earlier onset of tumorigenesis. To longitudinally assess liver cancer growth, we developed a novel *in vivo* method, utilizing 800 CW 2-deoxyglugose (800 CW 2-DG), to track disease progression in live animals by exploiting the elevated rate of glycolysis in tumour cells, even in anaerobic conditions (Rajendran et al., 2004). Using this approach, we were able to initially observe that at 28 weeks after DEN administration, the fluorescent signal in the *Clspn*^+/-^ livers was higher than in the WTs suggesting earlier onset of tumorigenesis (Fig. 7B). This was confirmed at 30 weeks, when the mice were imaged again (Fig. 7B). Following imaging, the mice were euthanised and macroscopic liver tumours were counted and also imaged *ex vivo*. This confirmed that the *Clspn*^+/-^ mice had significantly higher fluorescent signal in the livers *ex vivo* (Fig. 7C) and this corresponded to a significantly increased number of visible liver tumours (Fig.7D & E, S7A).

**Figure 7:**
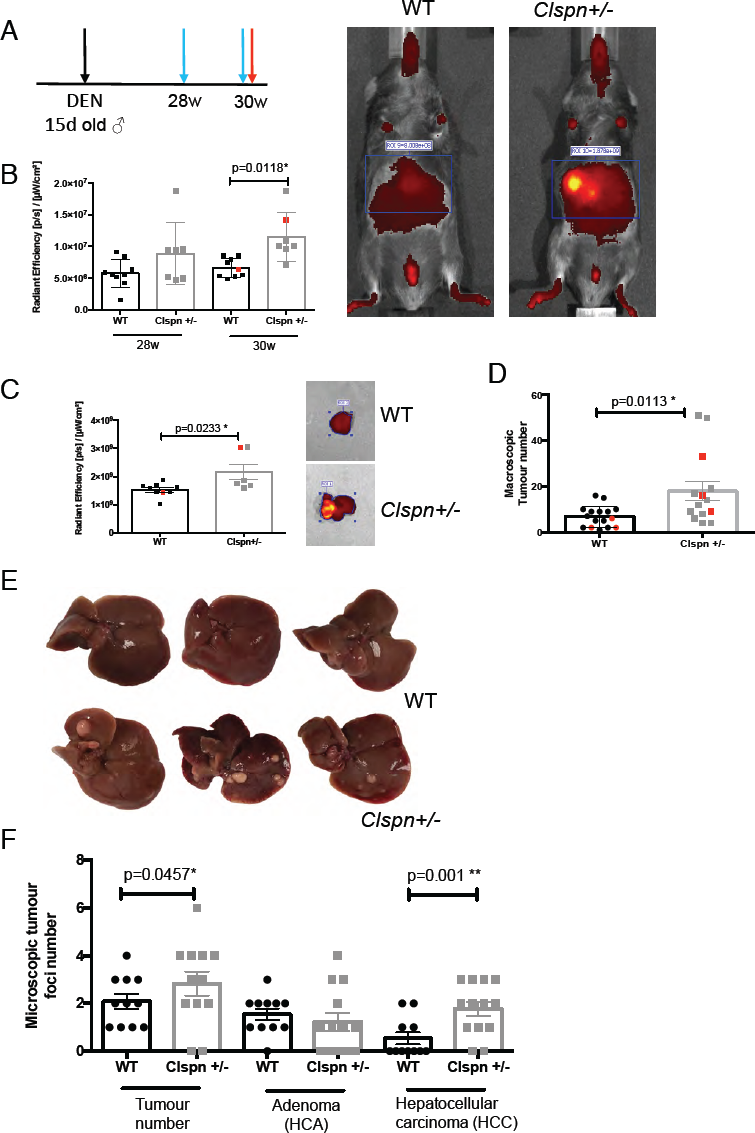
Earlier onset of DEN induced hepatocellular carcinoma in *Clspn*^+/-^ mice. (A) Schematic diagram illustrating the chronic DEN *in vivo* study in WT and *Clspn*^+/-^ mice. 15 day old mice were given 30 mg/kg DEN in 0.9% saline by IP injection. Black arrow indicates DEN administration, blue arrows indicate imaging timepoints and the red arrow indicates the time of study termination. (B) In vivo imaging using fluorescent 2-DG reveals increased tumour growth in *Clspn*^+/-^ mice. Quantification of the *in vivo* fluorescent signal from the abdominal region of either WT or *Clspn*^+/-^ mice either 28 w or 30 w post DEN administration. Data represents mean ± SEM. p<0.05 * (Unpaired Student’s t-test). Representative images are shown with red dots in scatter plots indicating representative samples chosen. (C) Quantification of the *ex vivo* fluorescent signal from the livers of either WT or *Clspn*^+/-^ mice excised 30 w post DEN administration. Data represents mean ± SEM. p<0.05 * (Unpaired Student’s t-test) Representative images are shown with red dots in scatter plots indicating representative samples chosen. (D & E) Livers from *Clspn*^+/-^ mice show increased numbers of visible tumours. Numbers of macroscopic tumours visible in WT and *Clspn*^+/-^ livers 30 weeks after DEN administration were counted (D). Data represents mean ± SEM. p<0.05 * (Unpaired Student’s t-test) Representative images (E) of livers in WT and *Clspn*^+/-^ livers 30 weeks after DEN administration. Red dots in (D) scatter plots indicate liver images chosen. (F) DEN treated *Clspn*^+/-^ mice have increased numbers of hepatocellular carcinoma. Histological quantification of tumours and classified as either hepatocellular adenomas or carcinomas in liver sections from DEN-treated WT and *Clspn*^+/-^ mice. An expert pathologist scored the pathology. Data represents mean ± SEM. p<0.05*, p<0.01** (Unpaired Student’s t-test).

Detailed histological analysis of liver tissue sections confirmed the visible differences in tumour numbers between *Clspn*^+/-^ mice and WT mice (Fig. 7F, S7B). Interestingly, while the numbers of adenomas are broadly equivalent between the two mice strains (17 foci from 11 WT mice, 16 foci from 13 *Clspn*^+/-^ mice), the more malignant hepatocellular carcinoma numbers were dramatically different. Here the 13 *Clspn*^+/-^ mice had 23 hepatocellular carcinomas, including one grade II HCC, versus only six seen in the WT mice (Figs. 7F, S9B). This result was strikingly similar to that we had previously reported in *RelA*^T505A^ mice (Moles et al., 2016), indicating that NF-κB regulation of Claspin levels provides an explanation, at least in part, for the ability of this transcription factor to regulate onset and progression of DEN-induced HCC.

## Discussion

It has been known for some time that DNA damage activated checkpoint kinases can regulate NF-κB, either through activation or inhibition of IκB kinase activity or direct phosphorylation of NF-κB subunits (Campbell et al., 2004; Hinz et al., 2010; Rocha et al., 2005; Sabatel et al., 2012; Schmitt et al., 2011; Wu and Miyamoto, 2008; Wu et al., 2010). However, it is less well established whether NF-κB itself regulates the DNA damage response and if this has any role in tumorigenesis. Previously, the Rocha lab demonstrated that in a variety of human and mouse cell lines, IKKβ and c-Rel regulate the *CLSPN* gene independently of the cell cycle (Kenneth et al., 2010). As a consequence they observed reduced CHK1 activity in response to ultraviolet light and hydroxyurea in which IKK/c-Rel activity was inhibited. However, this study did not examine the significance of this pathway *in vivo* nor any potential role in tumorigenesis. Here we have both confirmed and significantly extended these findings. We demonstrate that in two distinct mouse models, the *RelA*^T505A^ knockin mouse and the *c-Rel^-/-^* mouse, there is significant loss of Claspin expression and CHK1 activity and that this is associated with earlier onset of Eµ-Myc lymphoma. Low levels of *CLSPN* expression are also associated with reduced survival in some human forms of cancer (Fig. 5B, S5E, Table S2). Supporting this hypothesis we find that heterozygous knockout *Clspn* mice spontaneously develop B-cell lymphoma and treatment of these mice with the DEN carcinogen leads to earlier, more aggressive HCC. We propose that in the early stages of tumorigenesis, *CLSPN* is an NF-κB target gene that through promoting checkpoint kinase activation helps prevent further genomic instability and cancer development resulting from DNA replication stress (Fig. 8).

**Figure 8.**
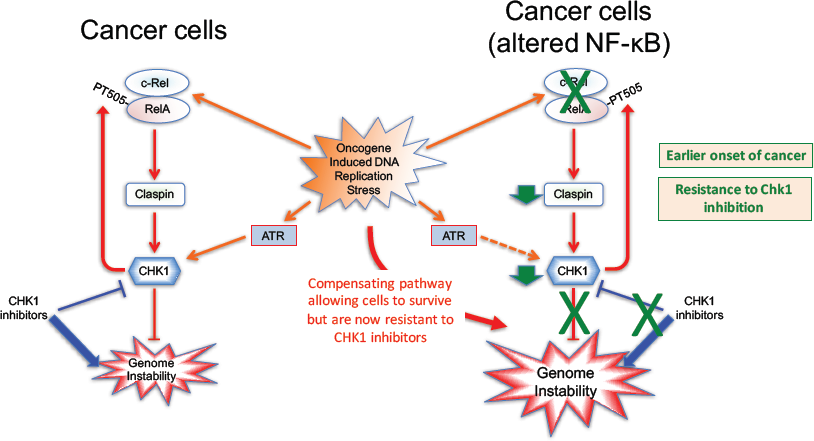
Schematic model illustrating NF-κB regulation of Claspin expression in cancer. Our data describe a positive feedback loop in which CHK1 phosphorylation of RelA at T505, together with the NF-κB subunit c-Rel, drives the expression of the ATR checkpoint kinase regulator Claspin in response to DNA replication stress in cancer cells. This in turn is required for maintenance of CHK1 activity. We propose that loss of this pathway early in tumorigenesis promotes cancer development through increased genomic instability. However, in malignant cancer cells it can help promote their addiction to the checkpoint kinase signalling required for the maintenance of genomic integrity. Importantly, disruption of this pathway leads to resistance of cells to treatment with CHK1 inhibitors.

Our results also imply a distinct role for NF-κB in the context of c-Myc driven cancer. Much of the research in the role of NF-κB in cancer has focussed either on its role in inflammation or haematological malignancies where activating mutations often lead to aberrant activity. However, there has been little analysis of its role in other types of the disease, such as c-Myc-driven lymphomas, with a normal, unmutated NF-κB pathway. We recently investigated the role of the c-Rel NF-κB subunit in the Eµ-Myc model (Hunter et al., 2016). Here we found that surprisingly and in contrast to the expected result, Eµ-Myc*/c-Rel^-/-^* mice displayed earlier onset of lymphoma that correlated with loss of the expression of the B-cell tumour suppressor BTB and CNC homology 2 (Bach2) (Hunter et al., 2016). Moreover, tumour suppressor-like activity has also been described for RelA in Myc driven lymphoma (Chien et al., 2011). Here, shRNA depletion of RelA was found not to affect progression of established lymphomas but did result in resistance to cyclophosphamide treatment due to an impaired induction of cell senescence (Chien et al., 2011). Similarly, inhibition of NF-κB through expression of a degradation resistant form of the inhibitor IκBα, revealed a requirement for NF-κB activity for therapy induced senescence in the Eµ-Myc mouse model of B-cell lymphoma (Jing et al., 2011). c-Myc can also inhibit expression of NF-κB2 (p100/p52), and loss of this NF-κB subunit in the Eµ-Myc mouse model resulted in impaired apoptosis and moderately earlier onset of disease (Keller et al., 2010). By contrast, deletion of NF-κB1 did not affect Eµ-Myc lymphoma development (Keller et al., 2005). These results imply a more complicated role for NF-κB subunits in Myc driven lymphoma than the tumour promoting activities usually associated with these transcription factors.

Although early clinical trials for CHK1 inhibitors have encountered problems these appear to result from off-target effects of these compounds (Carrassa and Damia, 2017) and newer more specific compounds entering clinical trials are not anticipated to encounter these issues (Blackwood et al., 2013; McNeely et al., 2014). Our results suggest that Claspin mRNA or protein levels could act as a biomarker for the use of CHK1 inhibitors in the clinic, although to date we have been unable to identify an anti-human Claspin antibody with sufficient specificity. Although *CLSPN* mutations are relatively rare in human patients they have been reported (Erkko et al., 2008; Zhang et al., 2009) and may also serve as a potential biomarker, but as yet this has not been investigated. Our studies have therefore identified a potential pathway through which tumours might become resistant to CHK1 inhibitors, either through reduced Claspin expression, mutations within the *CLSPN* gene and other regulators of CHK1 activity or within the NF-κB subunits that regulate their expression. These results also imply that in cells with low Claspin and CHK1 activity and displaying CHK1 inhibitor resistance, an alternative cell signalling pathway has become activated to allow the tumour to survive the on going genomic instability caused by DNA replication stress. Future studies identifying this pathway could lead to therapeutic strategies for patients developing resistance to CHK1 inhibitors.

Our analysis showed that in human breast, gastric and other cancer samples, low *CLSPN* mRNA expression correlated with reduced survival (Fig. 5B, S5E, Table S2). It is interesting to note that studies have shown that following double strand breaks Claspin can associate with BRCA1, a known breast cancer tumour suppressor, and is required for its phosphorylation by ATR (Lin et al., 2004; Yoo et al., 2006). Another report also suggested that high Claspin protein expression through stabilization by the deubiquitinase, USP20, correlated with better prognosis in gastric cancer (Wang et al., 2017). However, in other tumour types our analysis indicated the opposite effect, with low levels of *CLSPN* expression being associated with better survival (Table S2). These contrasting effects can be explained by the many roles Claspin has in DNA repair pathways that are distinct from its role in DNA replication. For example, Claspin is known to have a role in homologous recombination (Lin et al., 2004), mismatch repair (Liu et al., 2010) and nucleotide excision repair (Praetorius-Ibba et al., 2007), and therefore its role may depend on which pathway tumour cells are most dependent on, or become activated by, a given therapeutic regimen. Moreover, our data (Figs 5 & 8) suggest that Claspin loss or haplo-insufficiency is required at the point of tumour initiation in order to drive tumorigenesis but other tumours may downregulate, or fail to activate Claspin, at later time points as a consequence of tumour progression.

Taken together our data describe a previously unappreciated pathway through which NF-κB can function as a tumour suppressor and prevent cancer development. In this model, the NF-κB subunits RelA and c-Rel, in a manner dependent upon RelA T505 phosphorylation, drive the expression of Claspin in response to oncogene induced DNA replication stress. This in turn drives CHK1 activity, which early in tumorigenesis will inhibit cancer development through preventing the genomic instability required for additional genetic mutations. However, in malignant tumours, this pathway can help promote their addiction to the checkpoint kinase signalling required for the maintenance of genomic integrity and cancer cell survival. In our study, disruption of this pathway leads both to the earlier onset of cancer and to CHK1 inhibitor resistance. These results have important implications in the development of resistance in cancer patients treated with CHK1 inhibitors: they could lead both to the identification of cohorts likely to respond better to drug treatment as well as the development of alternative therapies in low responders and patients where resistance has developed.

## Methods

### Ethics statement

All mouse experiments were approved by Newcastle University’s Animal Welfare and Ethical Review Board. All procedures, including the of breeding genetically modified mice, were carried out under project and personal licences approved by the Secretary of State for the Home Office, under the United Kingdom’s 1986 Animal (Scientific Procedures). Animals were bred in the Comparative Biology Centre, Newcastle University animal unit, according to the FELASA Guidelines.

### Mouse models

*RelA*^T505A^ knock in mice were generated by Taconic Artemis (Germany) using C57Bl/6 ES cells (Moles et al., 2016), *c-Rel^-/-^* mice were provided by Dr Fiona Oakley (Newcastle University). Eµ-Myc mice were purchased from The Jackson Laboratory, Maine, USA. C57Bl/6 mice used for re-implantation studies were purchased from Charles River (UK). Male Eµ-Myc transgenic mice that were used as breeding stock were omitted from the survival analysis. *Clspn^+/-^* mice were generated by the Knockout Mouse Project using C57Bl/6 ES cells, and obtained *via* RIKEN (Saitama, Japan). *Clspn^+/-^* mice were maintained on a pure C57Bl/6n background. In all experiments, the relevant pure C57Bl/6 (WT) strain was used as a control. No blinding of groups in mouse studies was performed. All mice were designated to an experimental group dependent on their genotype.

### Drugs and compounds

CCT244737 was synthesized as described (Walton et al., 2012). MK8776 was purchased from Stratech (Ely, UK). Etoposide was purchased from Scientific Lab Supplies (Nottingham, UK), SN38 and Mitomycin C were obtained from R&D systems (Abingdon, UK). All other compounds were purchased from Sigma Aldrich.

### Generation of Mouse Embryo Fibroblasts (MEFs)

Heterozygote *RelA*^T505A^ mice were bred to produce a mixture of WT and *RelA*^T505A^ homozygote littermates. MEFs were isolated as follows. Internal torso connective tissue from 13.5-day embryos was washed in sterile PBS and minced in 1x Trypsin (Invitrogen) for 15 min at 37°C. Following repeated pipetting to break up large tissue fragments, the cell pellet was re-suspended in DMEM (Lonza) supplemented with 20% Fetal Bovine Serum (FBS) (Gibco, Paisley, UK) and 50U/ml penicillin/streptomycin (Lonza), and incubated at 37°C in a 5% CO_2_ humidified atmosphere. Once cells reached 90% confluency, they were subcultured in 75cm^2^ flasks and considered as passage 1. Cells were then cultured following the standard 3T3 protocol (Todaro and Green, 1963). Cells were considered immortalised beyond passage 14, but not used in experiments beyond passage 25.

### Generation of Ear Fibroblasts

Ear Fibroblasts (EFs) were generated as previously described (Jurk et al., 2014). Briefly, ear punch biopsies (two per mouse) were transported and stored in DMEM containing 10% FBS on ice. Punches were washed three times with serum-free DMEM, finely cut and incubated for 2hrs at 37°C in 2mg/ml collagenase A (Sigma Aldrich) in DMEM. A single-cell suspension was obtained by repeated pipetting and passing through a 25G needle. Cells were centrifuged for 10 min at 95 xg and cultured in Advanced DMEM/F-12 (DMEM, Invitrogen) plus 10% FBS, 50U/ml penicillin/streptomycin (Lonza), 2 mM L-glutamine (Lonza) at 37°C in a 5% CO_2_ humidified atmosphere. Each isolate was derived from a separate mouse.

### Apoptosis assays

*RelA*^T505A^ or WT MEFs were grown in 96-well plates and treated for 24 h with the appropriate concentration of Cisplatin (4 μg/ml), Etoposide (15 μM), Hydroxyurea (0.5 mM), SN38 (5 μM) or Mitomycin C (1 μg/ml). Cells were then harvested, lysed and assayed for Caspase-3 activity using the CaspACE Assay System (Promega, Southampton, UK), according to manufacturer’s guidelines. Samples were normalized to their protein concentration using the Pierce BCA Protein Assay Kit (ThermoFisher Scientific, UK).

### Cell viability assays

Freshly isolated Eµ-Myc, Eµ-Myc/*RelA^T505A^* or Eµ-Myc/*c-Rel*^-/-^ tumour cells (5×10^5^ per well), immortalized WT or *RelA*^T505A^ MEFs (5×10^3^) or primary WT or *Clspn*^+/-^ ear fibroblasts (2.5×10^4^) per well were seeded into 96-well plates. Increasing concentrations of the novel CHK1 inhibitor, CCT244747 (ICR, Sutton, UK), or MK8776 (Stratech, Ely, UK) or solvent controls were added to three replicate wells. After 96hrs, viability was quantified using the PrestoBlue Cell Viability Reagent (ThermoFisher Scientific, UK), according to manufacturer’s instructions.

### Cell survival assays

Exponentially growing immortalized WT or *RelA*^T505A^ MEFs were treated for 24h with 1 μM or 5 μM CHK1 inhibitor, CCT244747 (ICR, Sutton, UK), or MK8776 (Stratech, Ely, UK) or solvent controls before re-seeding onto Petri dishes at known cell number (1000, 2500 or 5000 cells/dish). Colonies were fixed 14 days later with methanol:acetic acid (3:1) and stained with 0.4% (w/v) Crystal Violet. Cloning efficiencies were normalized to untreated controls.

### Gene expression analysis using quantitative real-time PCR

Total RNA was purified from Eµ-Myc, Eµ-Myc*/RelA^T505A^* Eµ-Myc/*c-Rel*^-/-^ tumour cell pellets using the PeqGold total RNA extraction kit (Peqlab). From appropriate liver samples, total RNA was extracted from snap-frozen tissues by homogenisation using Precellys 24 ceramic mix bead tubes (Stretton Scientific Ltd) in a Precellys 24 benchtop homogeniser (Stretton Scientific Ltd) at 6500 rpm for 30 s. Following this, samples were passed through Qiashredders (Qiagen, Crawley, UK) and RNA was purified using the Qiagen RNeasy mini kit according to manufacturer’s instructions.

RNA was measured for purity and concentration with the NanoDrop1000 (ThermoFisher Scientific) and reverse transcribed using the Quantitect Reverse transcription Kit (Qiagen) according to manufacturer’s instructions. Quantitative real-time PCR was performed on 20ng cDNA, in triplicate, using predesigned Quanititect Primer assays (Qiagen) to the following murine genes; *Clspn, Atr, Atrip, Rad17, Topbp1, Chek1.* These samples were run and analysed on a Rotor-gene Q system (Qiagen), using murine *Gapdh* primers as an internal control. All CT values were normalised to *Gapdh* levels.

### Caspase-3 assay

Both adherent and floating cells were harvested by centrifugation, and cell pellets were washed in ice-cold PBS. Caspase activity was analysed with the CaspasACE^™^ colorimetric assay kit (Promega) using 25 µg of protein extract following the manufacturer’s instructions. Protein quantification was undertaken using the bicinchoninic acid (BCA) protein assay (Thermo-scientific, Rockland, USA)Absorbance at 405nm is directly proportional to Caspase-3 specific activity.

### Western blotting

Whole cell extracts were prepared from Eµ-Myc, Eµ-Myc*/RelA^T505A^* or Eµ-Myc*/c-Rel^-/-^* tumour cell suspensions or extracted direct from snap frozen pieces of tumour or liver tissue. Snap frozen tumour or tissue was lysed in PhosphoSafe^™^ Extraction Reagent using the Precellys24 ceramic mix bead tubes (Stretton Scientific Ltd) in a Precellys®24 homogeniser (Stretton Scientific Ltd) at 6500 rpm for 30s, then extracted according to the PhosphoSafe^™^ Extraction Reagent manufacturer’s instructions. In the case of tumour cell suspensions, cell pellets were washed with ice-cold PBS, and lysed using PhosphoSafe^™^ Extraction Reagent (Merck-Millipore, Watford, UK), according to manufacturer’s protocols. Protein quantification was undertaken using the BCA protein assay, and samples resolved by standard denaturing SDS-PAGE gels. Samples were transferred onto PVDF membrane (Merck-Millipore) before being probed with the primary antibody. Horseradish peroxidase-conjugated secondary antibodies (anti-mouse; Sigma (UK), anti-rabbit; Sigma, UK) and enhanced chemiluminscence reagent (Thermo-scientific, UK) were used for detection.

### Immunohistochemistry

Formalin-fixed tumour or liver tissues were paraffin-embedded sections and serial sections cut by the Molecular Pathology Node, Cellular Pathology, Royal Victoria Infirmary, Newcastle-Upon-Tyne. H&E staining was also undertaken by the Molecular Pathology Node.

Formalin-fixed paraffin-embedded tumour and liver sections were dewaxed and hydrated. Endogenous peroxidase activity was blocked with hydrogen peroxide and antigen retrieval was achieved using 1 mM EDTA. Tissue was blocked using an Avidin/Biotin Blocking Kit (Vector Laboratories, Peterborough, UK) followed by 20% swine serum in PBS and then incubated with primary antibodies overnight at 4°C. The following day slides were washed and incubated with biotinylated swine anti-rabbit (Dako, UK) followed by Vectastain Elite ABC Reagent (Peterborough, UK). Antigens were visualised using DAB peroxidase substrate kit and counterstained with Mayer’s haematoxylin. Immuno-stained cells were imaged using a DFC310 FX microscope (Leica Microsystems) and the images blinded (coded) prior to analysis by an independent party. At least 5 images per tissue at x 10 magnification were analysed using brown/blue pixel intensity using Adobe Photoshop.

### Antibodies

Antibodies used were c-Rel (sc-71 Santa Cruz), c-Myc (sc-42 Santa Cruz), RelA (sc-372 Santa Cruz), RelB (4954 Cell Signaling), p105/p50 (ab7971 Abcam), p100/p52 (4882 Cell Signaling), β-Actin (A5441 Sigma), CHK1 (phospho S345) (2341 Cell Signaling), CHK1 (2360 Cell Signaling) RelA (phospho 536) (3031 Cell Signaling), active caspase 3 (9664 Cell Signaling), PCNA (ab64100), γH2AX (9718 Cell Signaling) and CD45R (ab18197).

Antibodies to the murine form of Claspin was generated by Moravian Biotechnologies. Anti-rabbit IgG (A6154Sigma and 7074 Cell Signaling) and anti-mouse IgG (A9044 Sigma) HRP-linked secondary antibodies were used for western blot detection.

### Eµ-Myc mice studies

Eµ-Myc*/RelA*^T505A^*^+/-^* offspring were generated by mating *T505A* female mice with Eµ-Myc male mice, further Eµ-Myc/*RelA*^T505A^ mice were generated by crossing Eµ-Myc*/T505A^+/-^* males with *T505A* female mice. Only group housed Eµ-Myc/*RelA*^T505A^ males were included as the cohort for this analysis to minimise any potential effects from environmental and endogenous estrogens. Eµ-Myc*/c-Rel^+/-^* offspring were generated by mating *c-Rel^-/-^* female mice with Eµ-Myc male mice, further Eµ-Myc*/c-Rel^-/-^* mice were generated by crossing Eµ-Myc*/c-Rel^+/-^* males with *c-Rel^-/-^* female mice. To perform survival analysis, Eµ-Myc transgenic mice were monitored daily and were sacrificed at pre-determined end-points, defined as the animal becoming moribund, losing bodyweight/condition and/or having palpable tumour burden at any lymphoid organ site.

Survival analysis (Kaplan Meier analysis) was carried out using GraphPad Prism (Version 5.0) and significance determined using the log-rank (Mantel-Cox) test. Moribund mice were necropsied and single cell suspensions were prepared from tumour-bearing organs. Briefly, lymph nodes, spleen or thymus were homogenised through a cell strainer, and single cell suspension collected in DMEM (Lonza) supplemented with 10% FBS, 5mM L-glutamine, 5mM sodium pyruvate, 1μM L-asparagine and 50μM β-mercaptoethanol (Sigma Aldrich). These cell suspensions were then used for downstream analyses or frozen in 90% FBS/10% DMSO for long-term storage and transplantation.

Technical note: This study does not include analysis of survival in Eµ-Myc/*Clspn*^+/-^ mice. We were advised that the Director of the Newcastle University Faculty of Medical Sciences Comparative Biology Centre had serious ethical and 3Rs concerns with this option, due to the number of mice required to generate a sufficiently powered cohort of male Eµ- Myc/*Clspn*^+/-^ mice. Female *Clspn*^+/-^ mice have reduced litter sizes (2 versus 5 for wild type mice in 100 days), lengthened time to first litter (44.5 days versus 22 days for wild type mice) and reduced average pups per litter (3.8 versus 7.2 for wild types). Eµ-Myc mice can only be bred as heterozygote transgenes (homozygous are non viable) and Eµ-Myc females cannot be used to carry litters due to the early onset of disease. Consequently only 1 in 16 of the *Cslpn*^+/-^ mice (female) x Het Eu-Myc mice (male) would be male and have the Eµ-Myc/*Clspn*^+/-^ genotype. We would therefore only obtain a suitable male mouse approximately once every four successful matings.

### Reimplantation studies

For tumour therapy studies, 2×10^6^ Eµ-Myc, Eµ-Myc/*RelA*^T505A^ or Eµ-Myc*/c-Rel^-/-^* tumour cells from male mice were transplanted intravenously (IV) via the lateral tail vein into 8-week old male C57BL/6 recipients. Mice were monitored daily using parameters such as their bodyweight and food and water consumption to assess disease progression. Mice were necropsied when they became moribund and the tumour burden assessed.

Oral administration of the CHK1 inhibitor, CCT244747, prepared as previously described (Lainchbury et al., 2012) (ICR, Sutton, UK), or vehicle control (65% PEG-400, 20% Tween-20, 10% H_2_O, 5% DMSO (all Sigma Aldrich)) was initiated when tumours became palpable (approximately 10 days after inoculation of Eµ-Myc or Eµ-Myc/*RelA*^T505A^ cells, and 20 days after inoculation of Eµ-Myc*/c-Rel^-/-^* cells). CCT244747 was given as a single agent, bolus dose (100 mg/kg p.o.) for 9 consecutive days. Lymphoid tumour burden and final tumour weights were measured at necropsy 24 hours after the final dose.

### *In-vivo* models of liver injury and hepatocellular cancer

80 mg/kg DEN in 0.9% saline was administered intraperitoneally (IP) to 8 week old male mice for the acute liver injury studies. In this case mice were humanely killed 24h or 48h post-DEN and tissues collected. In order to induce liver cancer, day 15 mice were given 30 mg/kg DEN in 0.9% saline by IP injection. For this model, *in vivo* imaging was performed at both 28 weeks and prior to harvest at 30 weeks. 20 nmol 800 CW 2-deoxy-D-glucose (Licor, UK) was administered IV *via* the lateral tail vein, 24h prior to imaging at 740/800nm on the IVIS Spectrum system (Perkin Elmer) under isoflurane anaesthesia. Mice were humanely killed, and liver tissue was harvested and imaged *ex vivo* at 30 weeks post DEN.

### ChIP-Seq

ChIP Seq data shown here was extracted from a previously published analysis of the EBV-transformed lymphoblastoid B-cell line (LCL) GM12878 using validated anti-RelA, RelB, cRel, p52 and p50 antibodies (Zhao et al., 2014). GM12878 are one of three ENCODE project Tier 1 cell lines. It is an original HapMap cell line used in many genetic studies including the 1000 Genomes Project and has a relatively normal karyotype. Reads from all ChIP-seq experiments were mapped to the hg19/GRCh37 build of the human genome using the UCSC genome browser. NF-κB ChIP-seq datasets have been published (gene expression omnibus, accession code GSE55105) (Zhao et al., 2014).

### γH2AX Immunostaining by flow cytometry

Eµ-Myc or Eµ-Myc/*RelA*^T505A^ tumour cells were washed in PBS and fixed with 4% formaldehyde before permeabilisation with 0.1% Triton. Cells were then stained with either γH2AX-FITC conjugated antibody (560445 BD Biosciences, UK) or IgG-FITC conjugated control antibody (557782 BD Biosciences, UK) before a 2hr incubation protected from the light. Cells were then resuspended in DAPI staining buffer and analysed using the FACS Canto III (BD Biosciences) and DIVA (BD Biosciences) software.

### Analysis of immune cell populations

Spleens, lymph nodes, blood and peritoneal fluids were isolated. For blood samples, red blood cell lysis was performed prior to cell acquisition using FACS lysis buffer (Becton Dickinson) according to manufacturer’s instructions. Murine tissue single cell suspensions were generated followed by incubation with anti-Fc receptor blocking antibody (anti-CD16/CD32 clone 2.4G2, BD Pharmingen). Distinction between live and apoptotic/necrotic cells was performed based on staining with LIVE/DEAD Aqua (Life Technologies). For cell surface marker detection cells were stained with a combination of FITC, PE, APC, PerCPCy5.5, APC-Cy7 conjugated monoclonal antibodies (BD Pharmigen). For detection of Foxp3, intracellular staining was performed using a Fix/Perm kit (eBiosciences). See Table S1 for additional antibody information. All Samples were acquired using a three laser FACS Canto II (BD Bioscience) and the data was subsequently analysed using FlowJo (version 9 or X; Treestar, USA).

### Statistical analysis

GraphPad Prism software (http://www.graphpad.com, V6.0) was used for statistical analysis. Except where stated in figure legends, Mantel-Cox and unpaired t-tests were used to calculate P values (P values of p<0.05 were considered significant).

#### Acknowledgments

We would like to thank Sonia Rocha, Derek Mann, Claire Richardson, Elaine Willmore and all members of the NDP laboratory for helpful advice and assistance. We also thank Drs. Luis Barrera and Hufeng Zhou for assistance with bioinformatics analysis and Dr Thomas Matthews (Institute of Cancer Research, London) for synthesis of CCT244747. JEH is funded by Cancer Research UK grant C1443/A22095 and has previously received funding from Leukemia Lymphoma Research grant 11022 and Cancer Research UK grant C1443/A12750. JAB, HS and AJM were funded by the Wellcome Trust grant 094409. The IVIS^®^ Spectrum was funded by Welcome Trust Equipment grant 087961. BEG is funded by US National Institutes of Health (grant AI137337) and by a Burroughs Wellcome Medical Scientist career award. IC and MDG receive funding from Cancer Research UK grant number C309/A11566.

## Author contributions

JEH: performed majority of experimental work. Contributed to design of experiments and manuscript writing.

JAB: performed experiments with WT and *RelA*^T505A^ MEFs, assisted with procedures involving Eµ-Myc mice

HS: assisted with procedures involving Eµ-Myc and CLSPN^+/-^ mice. Performed γH2AX flow cytometry experiments.

SL: assisted with procedures involving imaging and liver *in vivo* models.

AF & AMK: performed analysis of immune cells in *RelA*^T505A^ mice.

AJM, NSK & RTC: contributed to experimental work.

HDT: provided training and assisted with lymphoma re-implantation studies.

KJC: provided advice on working with Eµ-Myc mice, assisted with data analysis and manuscript writing.

BEG: performed analysis of ChIP-Seq data. Assisted with manuscript writing.

FO: assisted with liver in vivo models and performed pathological analysis of DEN treated tumours. Assisted with manuscript writing.

MDG and IC: provided CCT244747 CHK1 inhibitor and contributed to the design and analysis of experiments. Assisted with manuscript writing.

NDP: contributed to design of experiments and manuscript writing.

### Conflict of Interest Disclosures

IC and MDG are current or former employees of The Institute of Cancer Research, which has a commercial interest in CHK1 inhibitors. The other authors disclose no conflicts of interest.

## Supplementary Figure Legends

**Figure S1:**
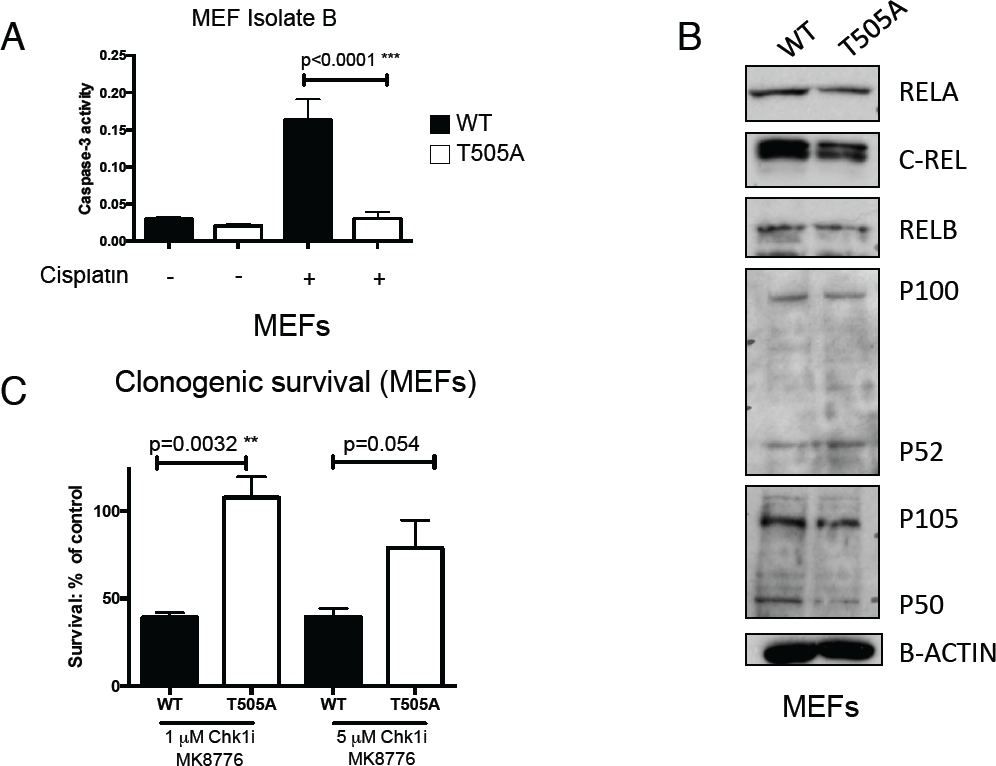
(A) *RelA*^T505A^ MEFs are resistant to Cisplatin induced apoptosis. CASPASE 3 activity assay in immortalised WT and *RelA*^T505A^ MEFs after 16 hours treatment with Cisplatin (4 μg/ml), Results shown are the mean + SEM from 3 separate repeat experiments. This is an independent isolate of the WT and *Rela^T505A^* MEFs to that used in Figure 1. (B) Western blot analysis of the NF-κB subunits, c-REL, RELA, RELB, p100/p52, p50 together with c-MYC in extracts prepared from WT and *Rela^T505A^* MEFs. (C) Increased clonogenic survival in *RelA*^T505A^ MEFs following CHK1 inhibitor treatment Clonogenic survival in WT and *RelA*^T505A^ MEFs following either treatment with either 1 μM (p=0.0032 ** Unpaired Student’s t-test) or 5 μM (p=0.0504 Unpaired Student’s t-test) of the CHK1 inhibitor, MK8776.

**Figure S2:**
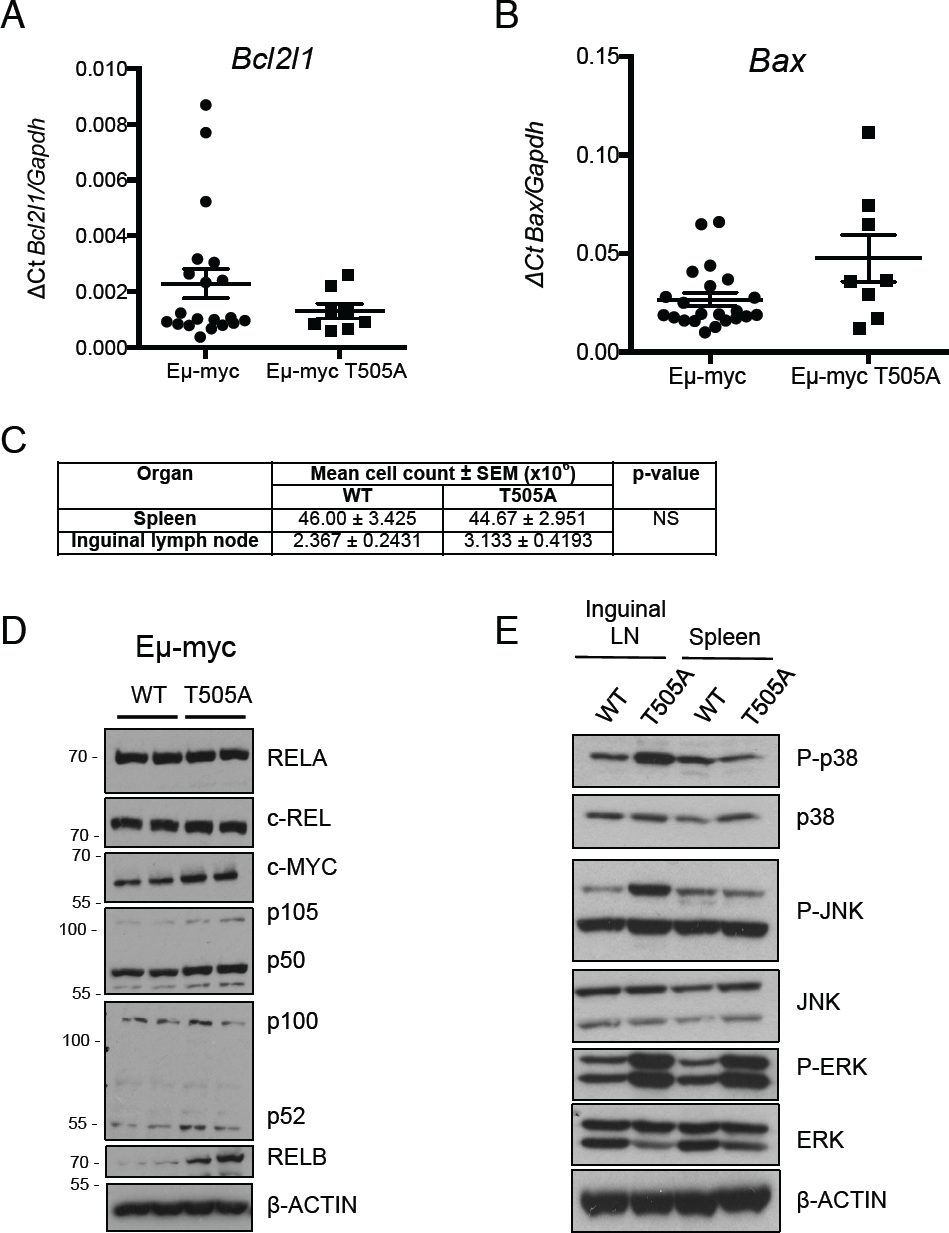
(A & B) There is no significant change in Bcl-xL (*Bcl2l1*) or *Bax* mRNA levels between Eµ-Myc and Eµ-Myc/*Rela^T505A^* lymphomas. qRT-PCR data showing relative *Bcl2l1* (A) *or Bax* (B) mRNA expression in tumorigenic spleens from Eµ-Myc and Eµ-Myc/*Rela^T505A^* mice. Data represents mean ± SEM, each point is an individual mouse. (C) Table detailing the cell numbers in spleen and inguinal lymph nodes from WT and *Rela^T505A^* mice. No significant difference was observed using a two-tailed Students t-test. (D) Western blot analysis of the NF-κB subunits, c-REL, RELA, RELB, p100/p52, p50 together with c-MYC in extracts prepared from Eµ-Myc and Eµ-Myc*/Rela^T505A^* mouse tumorigenic spleens. (E) Western blot analysis of the phospho p38 (Thr180, Tyr182), p38, phospho JNK (Thr183, Tyr185) JNK, phospho ERK (Thr202, Tyr204) or ERK in snap frozen tumours extracts prepared from Eµ-Myc and Eµ-Myc*/Rela^T505A^* mouse inguinal lymph nodes and spleens.

**Figure S3:**
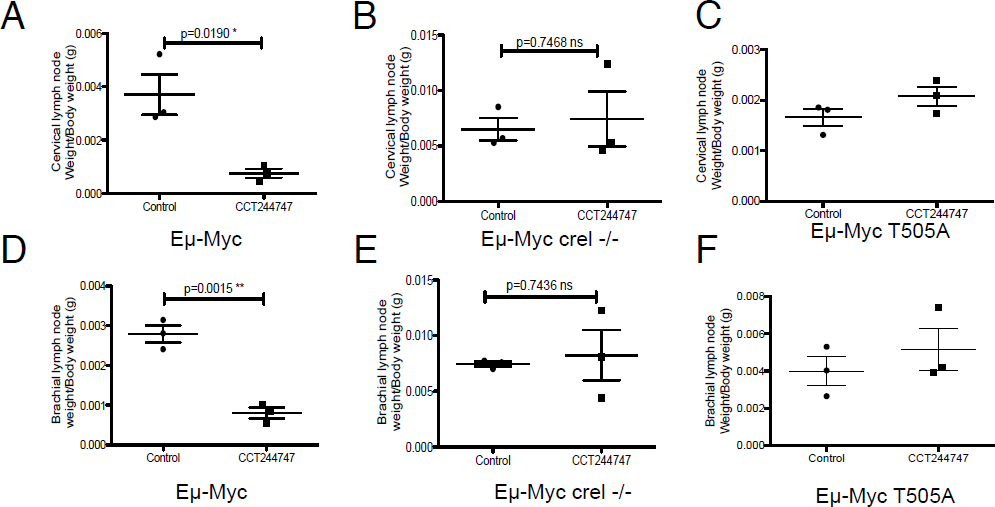
(A-F) Reimplanted Eµ-Myc/*c-Rel^-/-^ and* Eµ-Myc/*Rela^T505A^* lymphoma cells are resistant to CHK1 inhibition *in vivo.* Representative data from one Eµ-Myc or one Eµ-Myc/*c-Rel^-/-^* or one Eµ-Myc/*Rela^T505A^* tumour re-implanted into six C57Bl/6 mice and treated with CCT244747 (n=3), or vehicle control (n=3). Two organ sites are shown; cervical lymph nodes (A-C) and brachial lymph nodes (D-F), and in the case of all Eµ-Myc tumours, lymphoid tumour burden was significantly reduced by CCT244747 (data shown are mean ± SEM, p*<0.05, p**<0.01, p***<0.001 Unpaired Student’s t-tests). However, Eµ-Myc/*c-Rel^-/-^* or Eµ-Myc/*Rela^T505A^* lymphoid tumour burden was not significant affected by CCT244747 (data shown are mean ± SEM).

**Figure S4:**
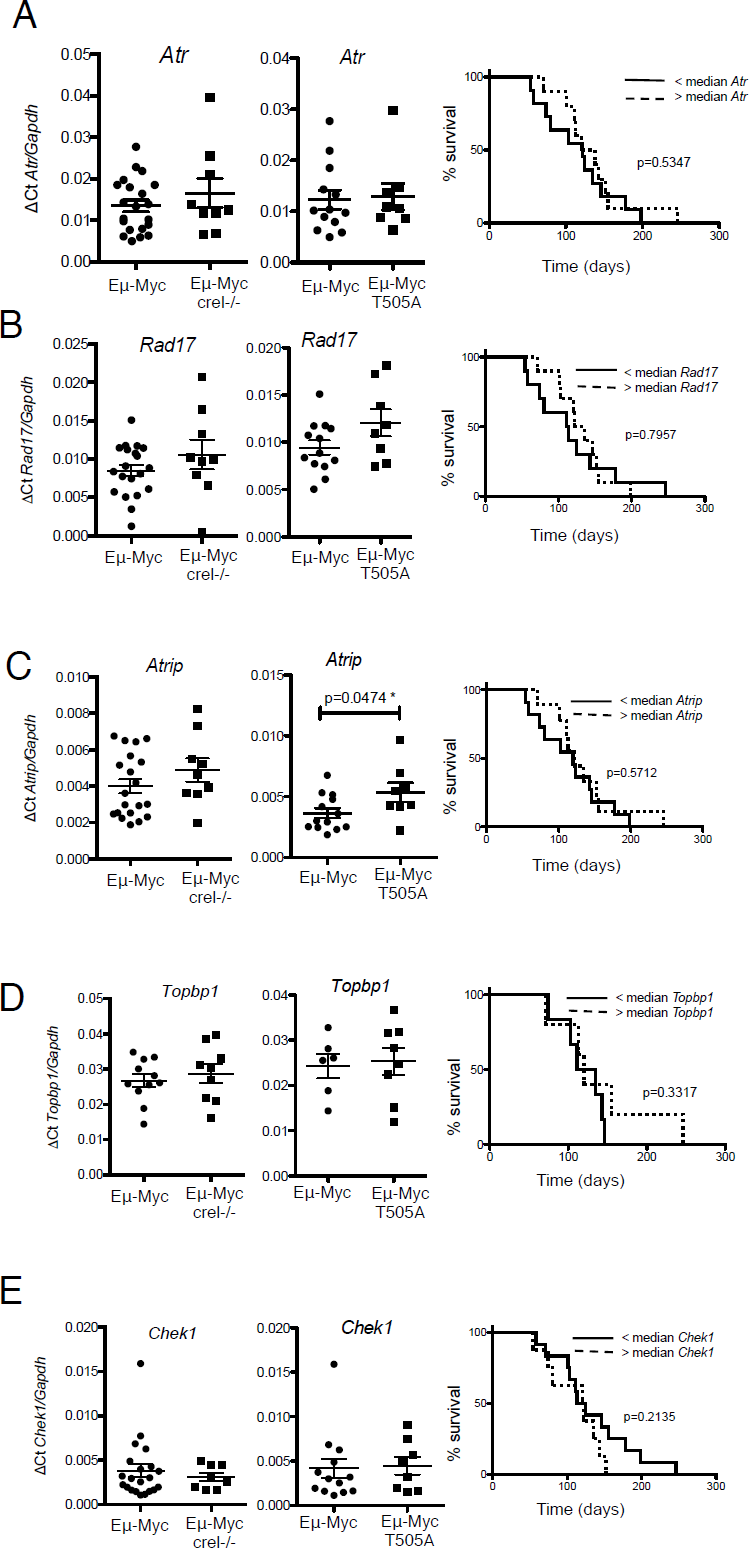
(A-E) Other components of the ATR/CHK1 signalling pathway are not affected in Eμ*-*Myc/*c-Rel^-/-^* and Eμ*-*Myc/*Rela^T505A^* mice and do not correlate with onset of lymphoma. qRT-PCR data showing relative *Atr* (A), *Rad17* (B), *Atrip* (C), *Topbp1* (D) and *Chek1* (E) expression in tumorigenic spleens from Eμ*-*Myc (for c-Rel analysis n=20 A-D, n=11 E, for *Rela^T505A^* analysis n=13 A-D, n=6 E), Eμ*-*Myc/*c-Rel^-/-^* (n=11) and Eμ*-*Myc/*Rela^T505A^* (n=8) mice. Right hand panel (A-E) shows Kaplan-Meier survival analysis of Eμ*-*Myc mice with below and above the median levels of *Atr, Rad17, Atrip, Topbp1* and *Chek1* mRNA (n=20 mice), and *Topbp1* (n=11 mice). No correlation with overall survival is seen.

**Figure S5:**
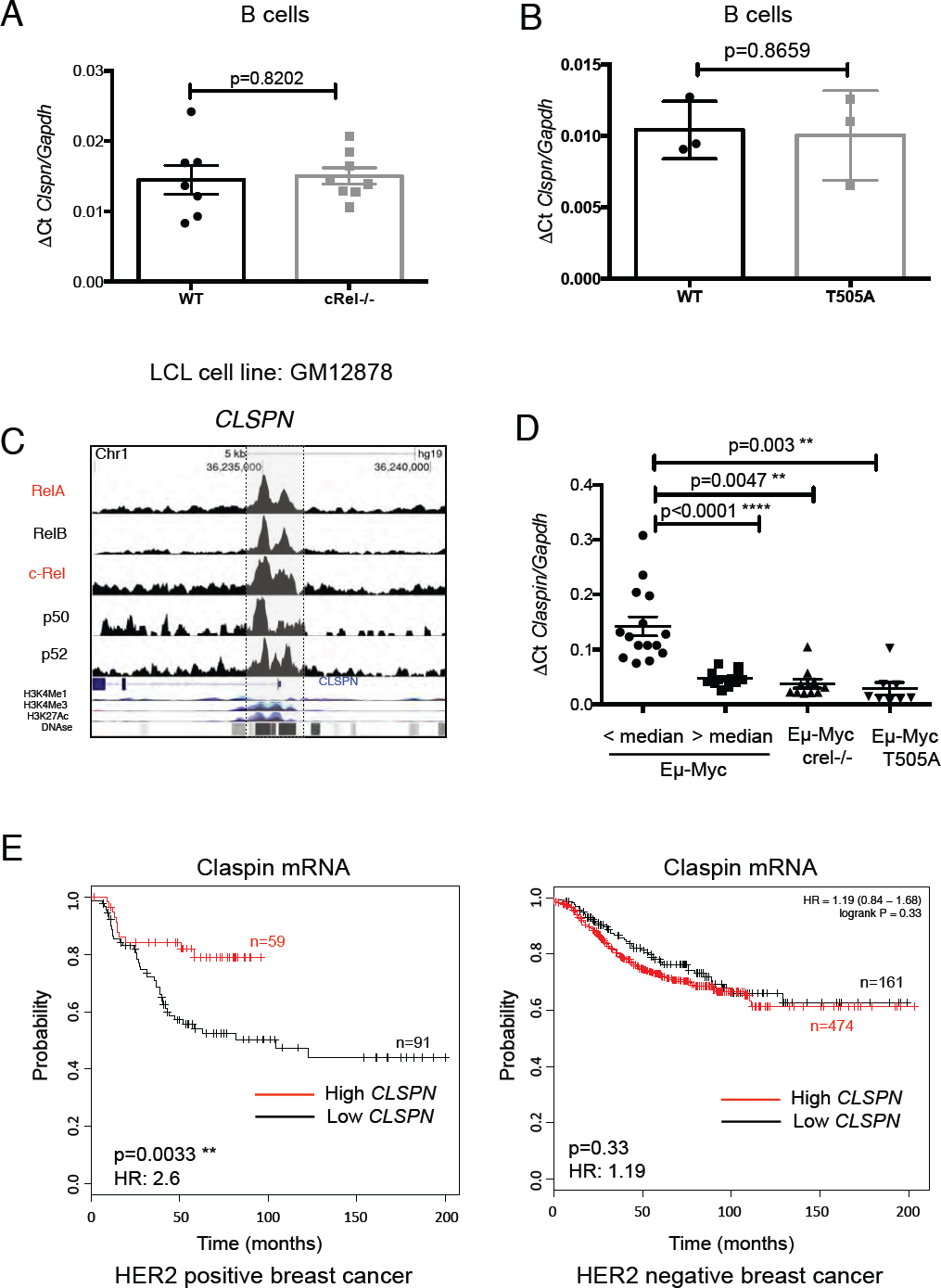
(A & B) *Clspn* mRNA levels are not affected in non-tumorigenic B-cells from *c-Rel^-/-^* (A) or *Rela^T505A^* (B) mice. qRT-PCR data showing relative *Clspn* expression in purified splenic B-cells from C57Bl/6 (n=6) and *crel^-/-^* (n=7) mice. Data represents mean ± SEM. Analysed using Unpaired Student’s t-test. (C) The *CLSPN* gene is a NF-κB target gene in B-cell lymphoma. ChIP Seq data showing NF-κB subunit binding to the human *CLSPN* gene promoter in the EBV-transformed lymphoblastoid B-cell line (LCL) GM12878. (D) Scatter plot showing how E*µ-*Myc mice were stratified into those expressing below or above the median levels of *Clspn* mRNA in tumours, as well as E*µ-*Myc*/c-Rel^-/-^* and E*µ-*Myc*/Rela^T505A^* mice. Analysed using an ONE-way Anova with Sidak post-hoc test (E) Low *CLSPN* mRNA levels are associated with poor survival in HER2 positive breast cancer patients. Kaplan-Meier survival analysis of breast cancer datasets from KM Plotter with differing levels of *CLSPN* mRNA (auto select best cutoff). Patients with low *CLSPN* levels have significantly shorter overall survival in HER2 positive patients (p=0.0033 Mantel-Cox test, HR 2.6). In HER2 negative breast cancer patients, *CLSPN* mRNA levels do not predict outcome in this context (p=0.33 Mantel-Cox test, HR 1.19).

**Figure S6:**
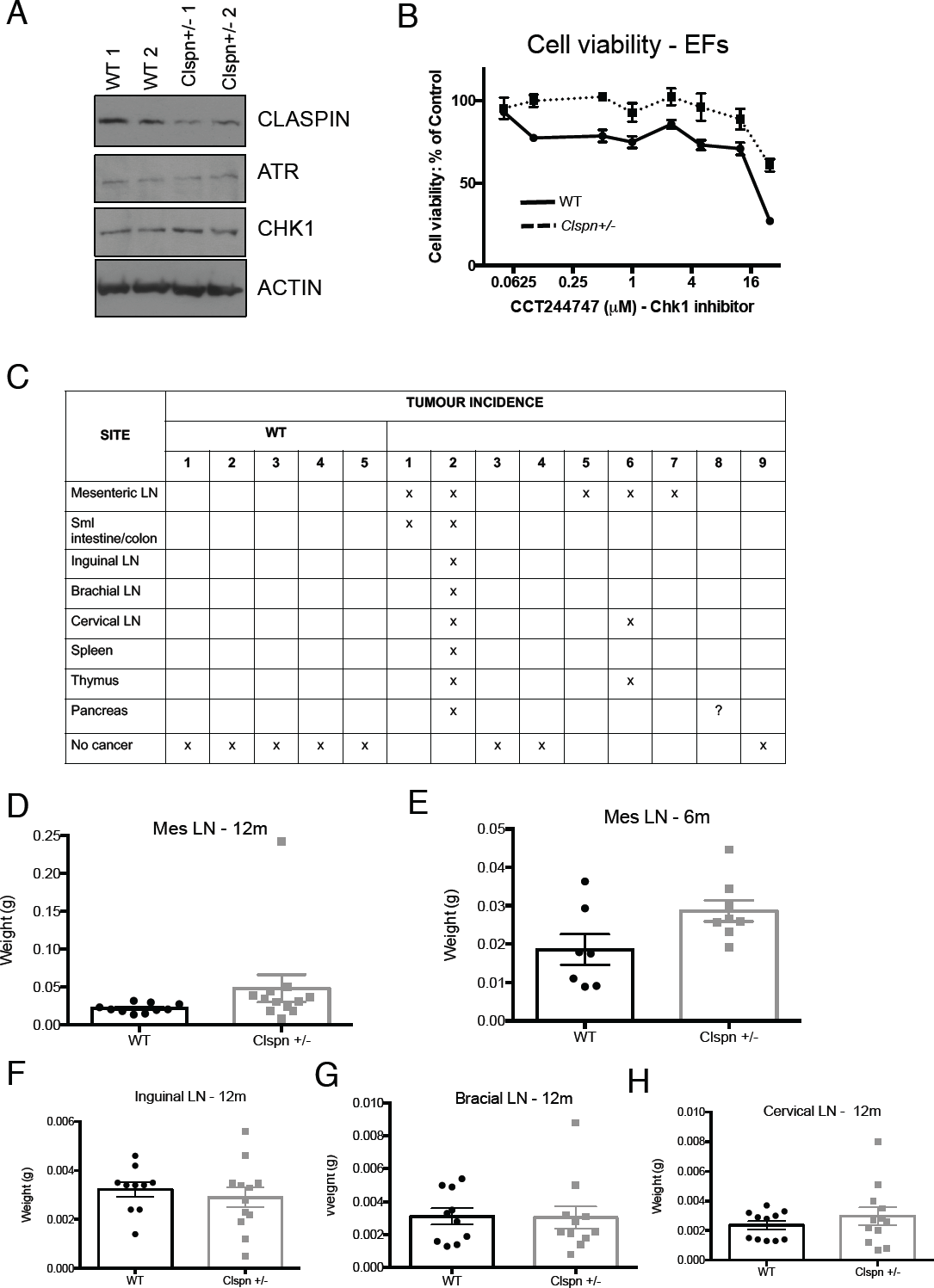
(A) Western blot analysis of Claspin, ATR and CHK1 in two independent isolates of WT and *Clspn*^+/-^ Ear Fibroblasts (EFs). (B) Primary fibroblasts from *Clspn*^+/-^ mice are resistant to CHK1 inhibition. Cell viability (Prestoblue assay) in WT and *Clspn*^+/-^ primary ear fibroblasts (EFs) following treatment with increasing concentrations of the CHK1 inhibitor, CCT244747. This is an independent isolate of EFs to those used in Figure 5C. (C) 18 month old *Clspn*^+/-^ spontaneously develop B-cell lymphoma. Table summarizing the tumour incidence in various organs in aged WT and *Clspn*^+/-^ mice. (D – H) Weight of the indicated lymph nodes from 12 month old (D, F-H) and 6 month old (E) WT and *Clspn*^+/-^ mice

**Figure S7:**
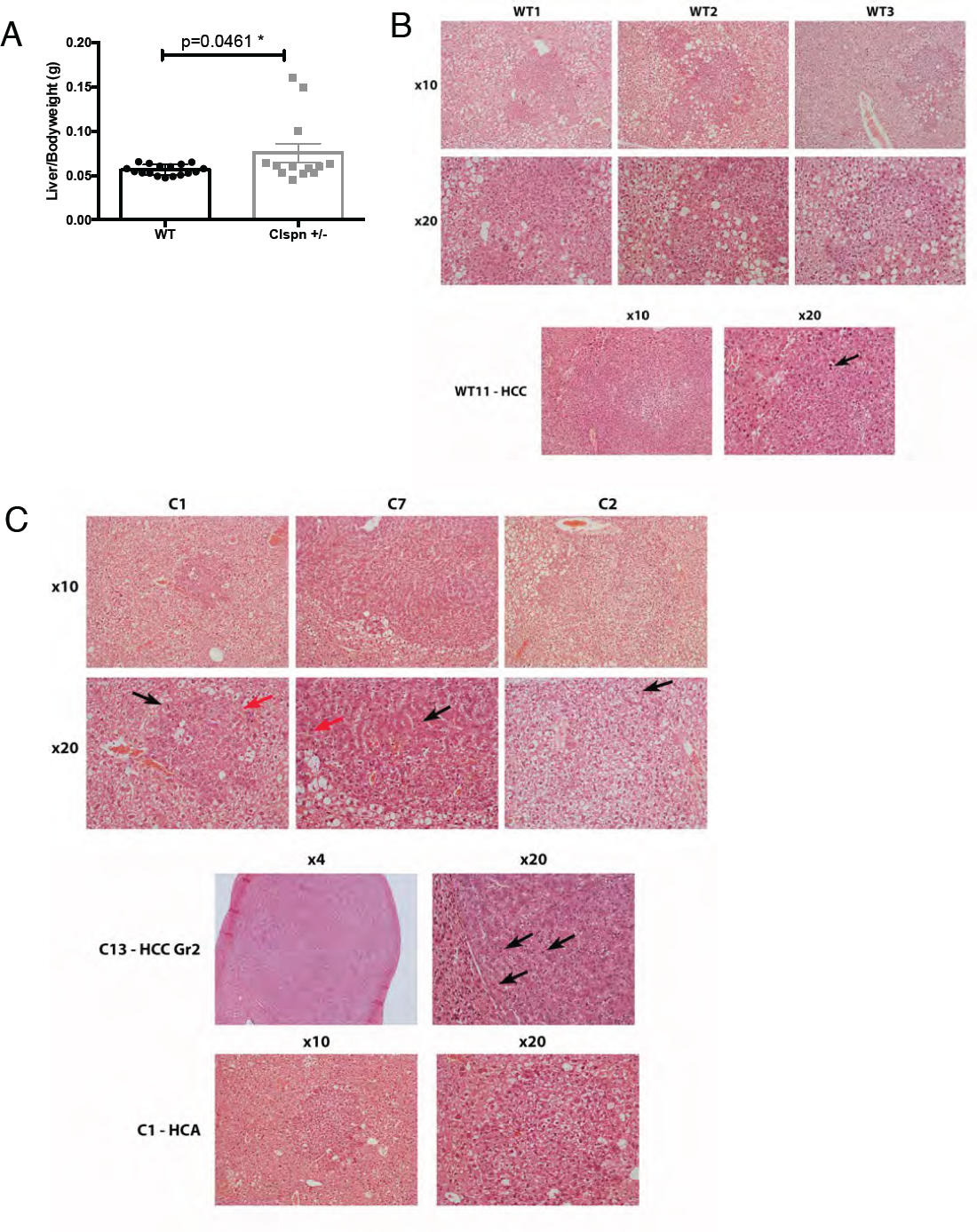
(A) Liver:body weight ratio at 30weks post DEN treatment in male WT and *Clspn*^+/-^ mice. Analysed using Unpaired Student’s t-test. (B) Representative IHC images from 30 week DEN treated WT animals. The images show the presence of adenomas but not hepatocellular carcinomas in these animals. (C) Representative IHC images from 30 week DEN treated *Clspn*^+/-^ animals. The images show the presence of hepatocellular carcinomas (HCC) in these animals, including a Grade II HCC. Red arrows denote the presence of proteoglycans and black arrows denote the presence of mitotic bodies.

## Supplementary Table 1

**Table S1.** Flow cytometric analysis of spleen, blood and inguinal lymph node cell populations of 6 x WT and 6 xT505A 12 week old male littermates. Peritoneal fluid control data is not shown. Data analysed using a two-tailed Students t-test.

| Cell marker(s) | Cell population | Spleen |  |  | Blood |  |  | Inguinal lymph nodes |  |  |
| --- | --- | --- | --- | --- | --- | --- | --- | --- | --- | --- |
|  |  | WT Mean<br>± SEM | T505A Mean<br>± SEM | p-value | WT Mean<br>± SEM | T505A Mean<br>± SEM | p-value | WT Mean<br>± SEM | T505A Mean<br>± SEM | p-value |
| CD19<sup>+</sup> | B cells as a % of total non-apoptotic cells | 46.12 ± 2.65 | 41.72 ± 4.15 | NS | 26.46 ± 8.76 | 27.94 ± 3.25 | NS | 20.28 ± 1.42 | 14.91 ± 3.53 | NS |
| CD3<sup>+</sup> | T cells as a % of total non-apoptotic cells | 40.75 ± 1.73 | 42.37 ± 3.59 | NS | 23.45 ± 2.90 | 25.08 ± 2.79 | NS | 75.25 ± 1.19 | 67.68 ± 3.13 | NS |
| CD138<sup>+</sup>/CD19<sup>+</sup> | Plasma cells as a % of CD19<sup>+</sup> cells | 23.73 ± 3.91 | 17.78 ± 2.94 | NS | 24.80 ± 4.00 | 18.82 ± 2.46 | NS | 33.60 ± 5.62 | 27.49 ± 3.06 | NS |
| CD11b<sup>+</sup>/CD19<sup>+</sup> | B1 B cells as a % of CD19<sup>+</sup> cells | 8.58 ± 1.43 | 9.12 ± 1.54 | NS | 34.07 ± 2.81 | 38.77 ± 5.30 | NS | 5.41 ± 0.46 | 6.22 ± 1.70 | NS |
| CD11b<sup>+</sup>CD19<sup>+</sup>CD5<sup>+</sup> | B1a B cells as a % of CD19<sup>+</sup>CD11b<sup>+</sup> cells | 19.90 ± 3.33 | 24.50 ± 6.03 | NS | 35.82 ± 3.95 | 35.52 ± 4.07 | NS | - | - | - |
| CD11b<sup>+</sup>/CD19<sup>+</sup>CD5<sup>-</sup> | B1b B cells as a % of CD19<sup>+</sup>CD11b<sup>+</sup> cells | 77.50 ± 4.01 | 73.33 ± 6.30 | NS | 64.23 ± 3.65 | 64.50 ± 3.69 | NS | - | - | - |
| CD4<sup>+</sup>/CD3<sup>+</sup> | CD4 T cells as % of CD3<sup>+</sup> cells | 60.17 ± 2.17 | 60.85 ± 0.67 | NS | 62.65 ± 4.33 | 64.98 ± 2.88 | NS | 52.93 ± 0.53 | 56.55 ± 1.95 | NS |
| CD8<sup>+</sup>/CD3<sup>+</sup> | CD8 T cells as % of CD3<sup>+</sup> cells | 32.88 ± 2.76 | 32.00 ± 1.08 | NS | 32.37 ± 4.59 | 31.30 ± 2.79 | NS | 45.15 ± 0.49 | 41.28 ± 1.83 | NS |
| CD3<sup>+</sup>/CD4<sup>+</sup>/CD25<sup>+</sup> | CD4<sup>+</sup>CD25<sup>+</sup> cells as a % of CD3<sup>+</sup> cells | 6.738 ± 0.59 | 6.19 ± 0.32 | NS | - | - | - | 4.64 ± 0.54 | 4.55 ± 0.55 | NS |
| CD3<sup>+</sup>/CD8<sup>+</sup>/CD25<sup>+</sup> | CD8<sup>+</sup>CD25<sup>+</sup> cells as a % of CD3<sup>+</sup> | 0.45 ± 0.085 | 0.44 ± 0.07 | NS | - | - | - | 0.34 ± 0.10 | 0.33 ± 0.06 | NS |
| CD4<sup>+</sup>/FOXP3<sup>+</sup> | T regs as % of CD4<sup>+</sup> T cells | 2.44 ± 0.24 | 2.62 ± 0.18 | NS | 0.37 ± 0.08 | 0.46 ± 0.10 | NS | 4.16 ± 0.14 | 4.17 ± 0.24 | NS |
| F4/80<sup>+</sup>CD11c<sup>-</sup> | Macrophages as a % of total no-apoptotic cells | 5.65 ± 2.42 | 7.41 ± 2.35 | NS | 2.97 ± 1.67 | 1.66 ± 0.81 | NS | 0.39 ± 0.14 | 0.29 ± 0.07 | NS |
| CD11c<sup>+</sup>F4/80<sup>+</sup> | Dendritic cells as a % of total non-apoptotic cells | 2.17 ± 0.34 | 2.07 ± 0.28 | NS | 2.34 ± 0.78 | 2.27 ± 0.99 | NS | 0.95 ± 0.05 | 0.80 ± 0.21 | NS |

**Table S2.**
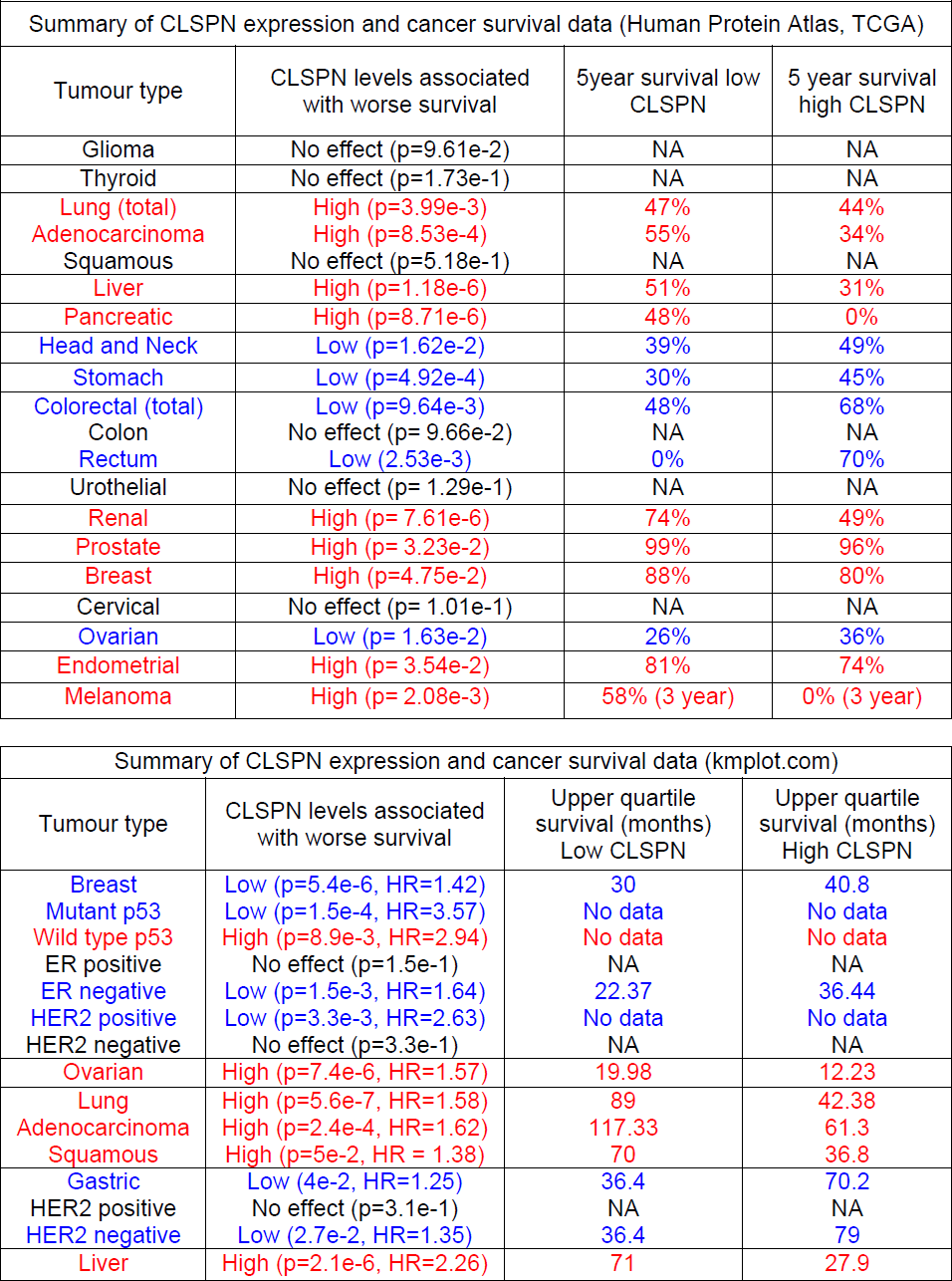
Summary of CLSPN mRNA expression and cancer patient survival data. Data analysed from TCGA datasets at Human Protein Atlas (www.proteinatlas.org) and KM plotter (kmplot.com). All data presented was using best separation cutoff. No effect is defined as p>0.05. For kmplot, data obtained from microarray analysis is with Jetset best probe selected (apart from Liver data which was obtained by RNA Seq).

